# CDC7 and CDK8 kinases cooperate to support DNA replication origin firing in human cells

**DOI:** 10.64898/2025.12.09.692982

**Authors:** Michael D. Rainey, Stefanus Bernard, Aikaterini Pravi, Liadhan Farrell, Dominik Boos, Colm J. Ryan, Corrado Santocanale

## Abstract

The coordinated activation of DNA replication origins is important for efficient DNA synthesis and genome stability. S-phase cyclin dependent kinases (CDKs) together with CDC7 kinase, are essential to origin activation by converting the pre-replicative complex into a fully active helicase.

To identify genes that tune DNA replication, we have performed a chemo-genetic genome-wide CRISPR-KO screen with cells challenged with the CDC7 inhibitor XL413. By developing a methodology based on genetic coessentiality to functionally cluster the hits, we uncover the transcriptional CDK8/CCNC kinase in a cluster with replication initiation factors. We find that CDK8 depletion further reduces the rate of DNA synthesis imposed by CDC7 inhibitors. DNA fibre experiments provide compelling evidence that CDK8 and CDC7 cooperate in origin activation.

Suppression of DNA synthesis by CDK8 inhibition requires the binding of CDK8/CCNC to the MDM2 Binding Protein (MTBP) and we show that CDC7 and CDK8 individually contribute to the phosphorylation of MCM4 subunit of the replicative helicase.

Thus, this work identifies CDK8 as the third protein kinase directly involved in origin activation in human cells.

## INTRODUCTION

In eukaryotic cells, DNA replication initiates from multiple origins (1–3). Two distinct steps facilitate initiation: the first is the loading of the core component of the replicative DNA helicase the MCM (Mini-Chromosome Maintenance) proteins at origin DNA to form pre-replicative complexes (pre-RCs), also described as origin licensing. The second step, origin firing, generates two bi-directional replisomes per pre-RC (4).

The current model involves cell-cycle cyclin-dependent kinases (CDKs) and the CDC7 kinase coordinating the molecular events at replication origins by phosphorylating several proteins of the pre-replicative complex. Phosphorylation of and association with additional factors result in the activation of the MCM helicase, thus leading to the local unwinding of double-stranded DNA at origins of replication. Unwinding is required for the loading of DNA polymerases and accessory factors that catalyse the semi-conservative synthesis of new DNA strands during chain elongation (5).

In budding yeast, CDC7 is essential for replication origin firing by phosphorylating multiple subunits of the MCM2-7 helicase complex facilitating the loading of CDC45 (6, 7). CDC7-dependent phosphorylation of the N-terminal tail of MCM4 was shown to counteract an autoinhibitory activity for MCM activation (8). Recent structural insights based on biochemically reconstituted DNA replication initiation showed that CDC7, together with its activating subunit DBF4 binds to the MCM2-7 double hexamer (DH-MCM), thus positioning its catalytic core in the proximity of the MCM4 and MCM2 terminal tails for subsequent phosphorylation (9–11).

In human cells, the CDC7 kinase activity is regulated by the binding of either DBF4 or DBF4B, also known as DRF1. The CDC7 gene and the kinase activity of its encoded product are absolutely required for cell proliferation in MCF10A breast cells (12). Instead DBF4 and DRF1 appear to be partially redundant for genome duplication and cell growth with DBF4 sustaining the bulk of CDC7 activity in MCF10A cells (13). At present, the molecular mechanisms of human MCM activation by CDC7 are not fully elucidated, but we and others have mapped specific CDC7-dependent phosphorylation sites on the MCM2 protein and showed that phosphorylation levels of MCM2 at Ser40/41 and Ser53 are fully CDC7-dependent. These phosphosites are good markers of cellular CDC7 activity and have proven robust pharmacodynamic markers in the discovery and development of CDC7 inhibitors (14–16). Furthermore, CDC7 dependent phosphorylation of hMCM4 was demonstrated by altered electrophoretic mobility in SDS-PAGE upon CDC7 inhibition (17), altogether suggesting that the molecular mechanisms of origin activation and the role of CDC7 kinase in origin activation is conserved throughout evolution (18).

CDC7-dependent phosphorylation of the MCM complex is counteracted by protein phosphatase 1 (PP1) which is targeted to the MCM complex by RIF1, thus the efficiency of initiation is dictated by a balance between phosphatase and kinase activity (19, 20). Importantly the RIF1-PP1 negative regulation of origin firing is reinforced by ATR-CHK1 signalling which, through the downregulation of CDK1 activity prevents the dissociation of PP1 from RIF1 (21). In addition to origin activation, CDC7 kinase has important roles in the replication stress response and chromatin function (reviewed in (7, 22)). Recent studies have shown that CDC7 plays a critical role directly at replication fork, promoting their restart after stalling and modulating fork speed (23, 24).

ATP-competitive CDC7 inhibitors (CDC7is) have been developed as an alternative strategy to restrain DNA replication by limiting origin firing instead of directly blocking fork progression (25). Inhibition of CDC7 leads to decreased origin activation, delayed S-phase progression and activation of the S-phase checkpoint, accompanied by mild replication stress and mitotic abnormalities (12, 16, 24). Replication stress involves ETAA1-dependent activation of ATR to further suppress origin activation and prevent premature entry and catastrophic progression through an aberrant mitosis with under-replicated DNA (26). The genetic background of the cells strongly contributes to the antiproliferative activity of CDC7i which varies substantially in different cell lines, and it is dependent on CDC7is potency in suppressing DNA synthesis and secondary responses driving either senescence or apoptosis (16, 23, 27, 28).

Alongside CDC7, cell cycle cyclin-dependent kinases control replication initiation. CDK kinases separate origin licensing, which is the loading of MCMs onto origins, from firing by inhibiting licensing and activating firing. cell cycle CDKs restrain licensing to the G1 phase when CDK activity is low, whilst enabling firing in S phase upon the rise of CDK activity. To inhibit licensing, CDKs directly phosphorylate MCM subunits (29–34) as well as other pre-RC components such as ORC and CDC6. To activate firing, CDKs phosphorylate Treslin/TICCR on two key residues, T969 and S1001, which is necessary for the interaction with TOPBP1 (35, 36). Treslin also binds to MTBP forming a trimeric complex (TOPBP1/Treslin/MTBP) (37) that facilitates the full assembly of the CDC45/MCM/GINS (CMG) complex and helicase activation. Studies in *Xenopus laevis* egg extracts indicate that CDC7 cooperate with CDKs in stabilising the TOPBP1/Treslin/MTBP complex (38).

To understand how the problems in DNA replication arising from CDC7 inhibition are dealt with, we performed a genome-wide CRISPR/Cas9 chemo-genetic screen directed to discover genes, that upon loss, could sensitize to the anti-proliferative effects of CDC7i. Among the hits we identified CDK8 and its cognate Cyclin, Cyclin C (CCNC).

CDK8 is a ubiquitously expressed member of the transcriptional CDK kinases. It is known as a component of the “Mediator-CDK8 module” (39, 40). The mediator of transcription, short Mediator, is a large multi-subunit complex which bridges transcriptional control elements, promoters, chromatin modifiers and transcription factors to RNA polymerase II (41, 42). The “CDK8 module” of the mediator is a four subunit regulatory subcomplex comprising CDK8, CCNC, MED12 and MED13. MED12 binding to CDK8-CCNC replaces the activation of the kinase by CAK-mediated t-loop phosphorylation that is common in other CDK kinases (43, 44). MED13 enables the interaction with the Mediator (40, 42). In mammals a paralog of CDK8, namely CDK19, exists and has high sequence similarity to CDK8. CDK19, in a mutually exclusive manner with CDK8, can form an analogous “CDK19 module” (45). Paralogs of MED12 and MED13 also exist, termed MED12L (MED12-like) and MED13L (MED13-like) respectively.

CDK8 is regarded as a proto-oncogene in colorectal cancer (CRC), as its high expression correlates with poor patient survival and with advanced disease (46, 47). Although CDK8 is generally not required for proliferation in most cell types (https://depmap.org/portal) (48), downregulation of CDK8 reduced proliferation of CRC cells *in vitro* and in xenograft models with CDK8 amplification, highlighting the therapeutic potential of targeting CDK8 (46, 47, 49). As a result, several CDK8 inhibitors have now been developed as targeted agents and, because of the similarity between CDK8 and CDK19 active site, most of these compounds can be considered as dual CDK8/19 inhibitors (42, 50, 51).

CDK8/CCNC can bind to MTBP, an essential DNA replication origin firing factor (37, 52, 53). This association occurs independently from MED12 and MED13, suggesting that this could represent a role independent from the Mediator complex and transcription (52). MTBP-CDK8/CCNC interaction is mediated by a metazoan specific binding region in MTBP that is important for efficient DNA synthesis, preventing replication stress and ensuring complete genome duplication to support a normal mitosis (52). MTBP binding to CDK8/CCNC, similarly to MED12 interaction, allosterically activates the kinase by repositioning the T-loop of the kinase independently of T-loop phosphorylation (54). However, while CDK8-CCNC can bind and phosphorylate MTBP *in vitro,* the relevance of CDK8 dependent phosphorylation of MTBP in living cells has not yet been assessed and more importantly a possible role of CDK8/CCNC in DNA replication has been elusive.

In this work we identify CDK8 as a main determinant for the antiproliferative activity of CDC7 inhibitors and we reveal for the first time a direct role of this kinase in the regulation of DNA replication cooperating with CDC7 to fire replication origins in human cells.

## RESULTS

### A genome-wide CRISPR/Cas9 loss of function screen identifies genes that affect proliferation rate in response to a CDC7 inhibitor

In order to identify genes that affect proliferation rate when DNA synthesis is partially restricted by mild CDC7 inhibition, we performed a CRISPR/Cas9 genome-wide knock-out screen (Fig. 1A). Briefly, MCF10A EditR cells that stably express Cas9 were transduced with a lentiviral-sgRNA library targeting approximately 19,000 coding genes and then were either untreated or treated with 1.25 µM of selective ATP competitive CDC7 inhibitor, XL413 (55). The concentration of XL413 used only mildly suppressed cell proliferation in this cell line (Supplemental Fig. S1A-C). CRISPR/Cas9-mediated loss of function of genes involved in supporting proliferation in untreated cells or in response to CDC7 inhibition is expected to result in a decreased relative abundance of the corresponding sgRNAs in the cell population. Next generation sequencing was then used to determine sgRNA abundance at day 0, after 20 days of growth in standard medium and after 24 days in media supplemented with XL413, when cells reached the same number as the control branch. sgRNA library representation was maintained throughout the experiment (Supplemental Fig. S1D).

**Figure 1:**
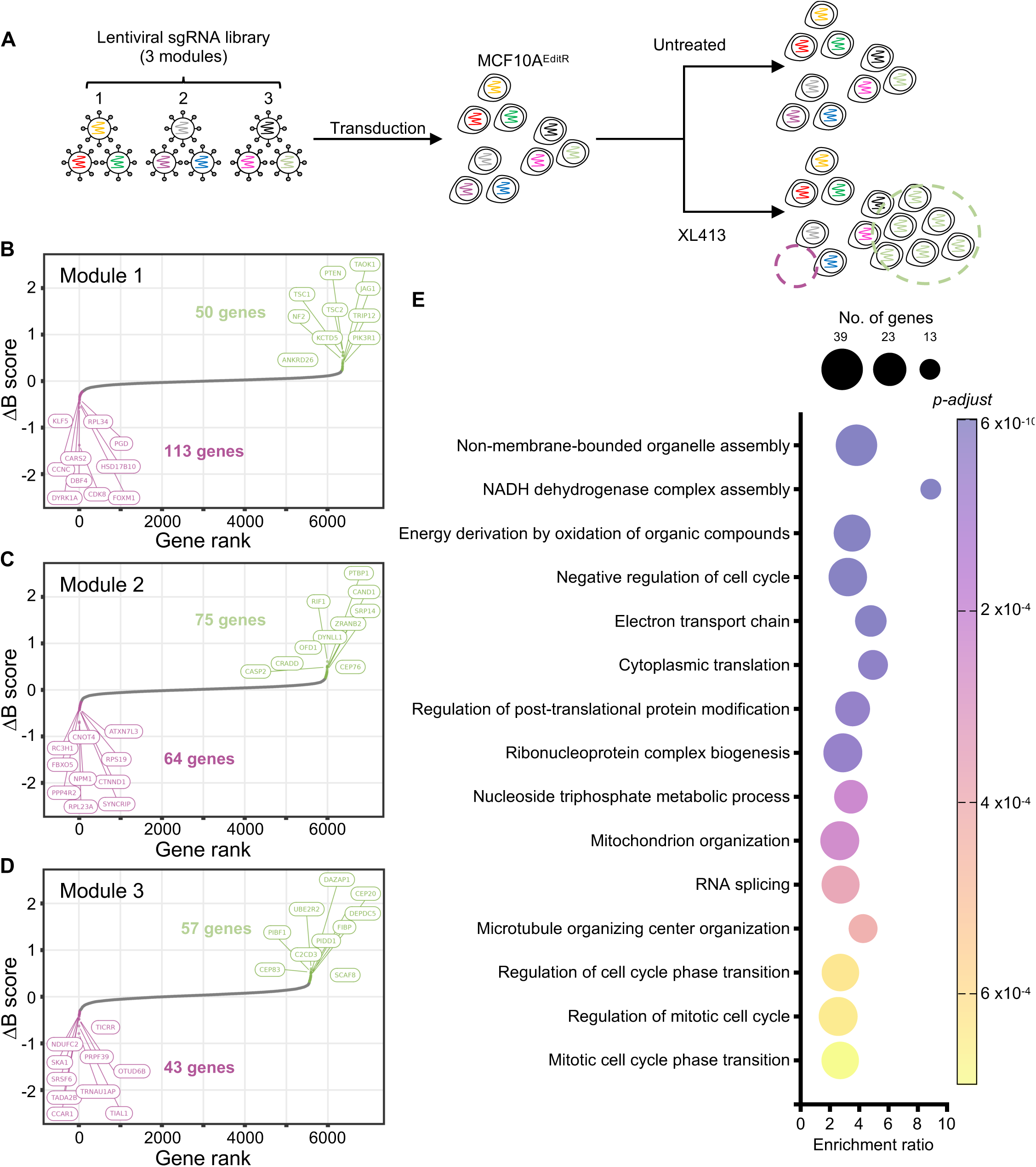
Genome-wide CRISPR/Cas9 screen with low doses of CDC7 inhibitor XL413. **(A)** Schematic representation of the screen’s workflow. MCF10 EditR were transduced with the three modules of the Cellecta library. After gene editing, cells were divided into two pools and cultured with or without 1.25 µM XL413. Clones with knock-out of genes conferring hypersensitivity are expected to be depleted in the treated population (magenta circles) while clones with knock-out of genes conferring resistance are overrepresented (green circles). **(B-D)** Plots showing genes ranked according to differential Beta score (ΔB score) between control and treated populations, for each module. Hits with lowest and highest ΔB score in each module are indicated in magenta and in green respectively and top 10 genes in each category are labelled. **(D)** Panel with the 15 most significant gene ontology terms, obtained by over-representation analysis of all the screen hits and a indication of the number of genes contributing to the enrichment ratio.

To further increase confidence in the screen we used a bioinformatic approach to flag and dismiss from consideration those sgRNAs in our library that perfectly aligned to more than one location within the GRCh38 human reference genome (56) and also those sgRNAs aligning in multiple locations with single and double mismatches in the PAM distal region, all of which had the strong potential of causing more than one dsDNA break in the genome (57) (Supplemental Fig. S1E).

The potential effects caused by individual gene KOs on fitness was independently determined through MAGeCKFlute pipeline (58) and output as a Beta score in both unchallenged and CDC7i treated populations. Altogether for each gene the combined result of the screen is shown as differential Beta score (ΔB score) between control and treated populations (Fig. 1B-D).

Utilizing a three standard deviation threshold as a cut-off, we identified 402 genes, 220 with negative ΔB score, which should include those that when knocked out, potentially decrease the proliferation rate specifically in CDC7i treated cells and 182 that, when knocked out could potentially relieve cells from the antiproliferative effect of the CDC7i. Among the latter we found *RIF1*, *ETAA1* and *PTPB1* which we had previously characterised, thus further giving confidence to pursue the analysis of the hits (Supplementary Table S1).

### Genetic co-essentiality describes *CDK8* and *CCNC* as DNA replication initiators factors

Gene Ontology analysis with all 402 hits indicates transport, mitochondrial respiration, protein translation and RNA metabolisms and cell cycle regulation (Fig. 1E) as processes involved in the cellular responses to CDC7 inhibition.

To understand how these genes could work together we developed a method to visualize genetic interaction by using gene co-essentiality analysis (59). Gene co-essentiality is a method to identify pairs of genes whose loss causes similar effects on fitness across a panel of cancer cell lines (60). Often pairs of genes that function in the same pathway or complex are essential in the same sets of cell lines and thus co-essentiality can reveal, in an unbiased fashion, functional relationship between genes. We performed gene co-essentiality analysis on genome-wide CRISPR-KO screens in 1,178 cell lines from DepMap 2024 Q4 (see methods) to identify gene pairs that were co-essential across this diverse set of cell lines, identifying a network containing a total of 25,133 co-essential gene pairs. We then integrated the hits from our screen with this network to identify the functional relationships between gene that caused resistance or sensitivity to CDC7i. In total, 249 interactions were identified between 165 hit genes from our screen, as presented in Figure 2. This network was further deconvoluted into 19 subnetworks (connected components with ≥ 3 genes, see methods) which were analysed for Gene Ontology enrichment. This revealed 15 subnetworks that were enriched in specific Gene Ontology terms including DNA replication, DNA repair, cilium organization (Fig. 2).

**Figure 2:**
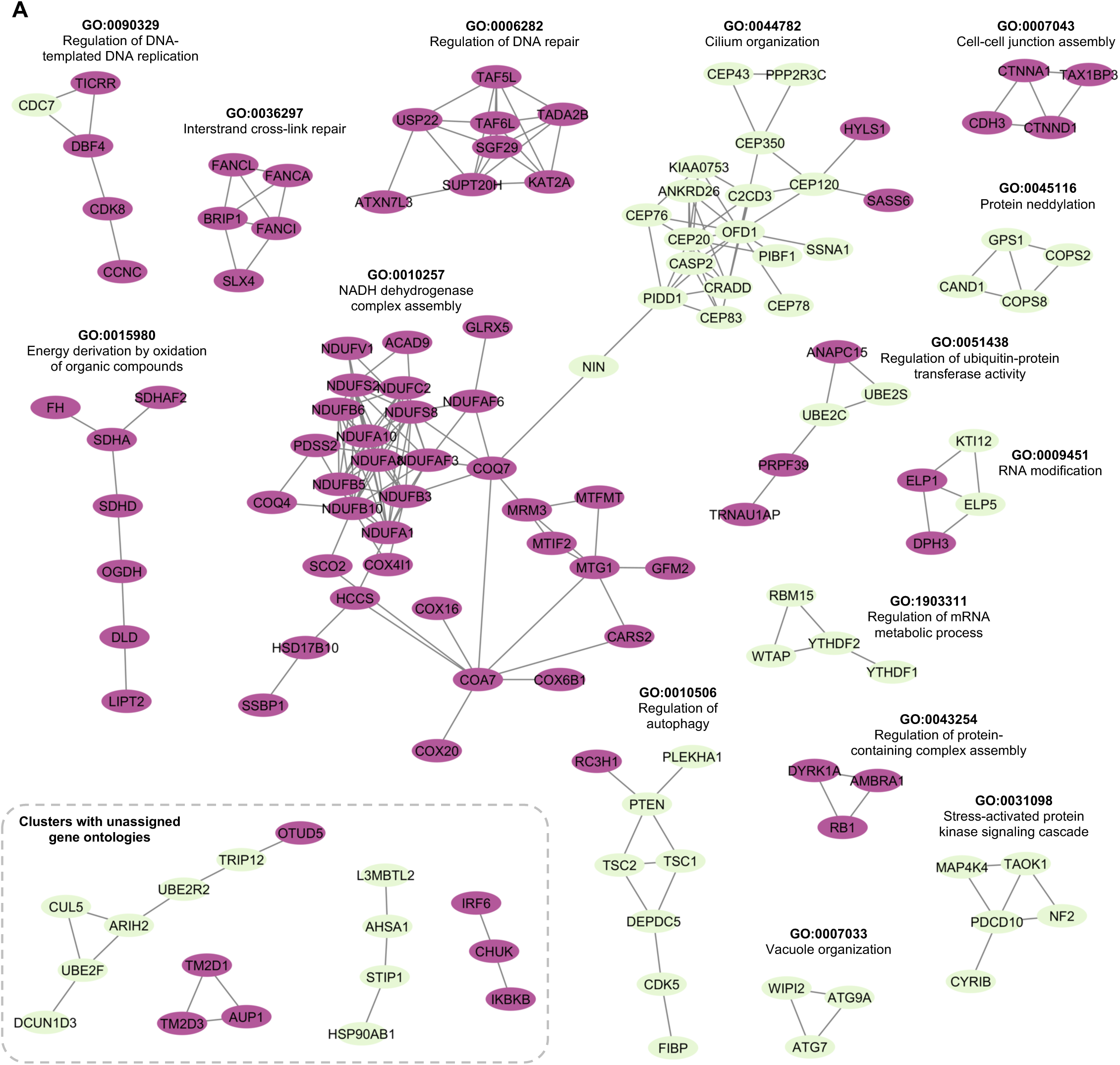
Co-essential genetic interactions among potential modulators of CDC7i anti-proliferative activity. **(A)** Co-essential genetic interaction networks among gene hits. Genes that when knocked-out are associated with negative ΔB score, thus potentially enhancing the anti-proliferative effects of CDC7 are in magenta, while the ones associated with positive ΔB score are in green. Gene ontology term and identifier are shown alongside associated subnetwork. Each edge indicates a significant co-essential relationship between a pair of genes (p-value < 0.05, FDR < 0.1) indicating that the two gene tend to cause correlated fitness defects when viewed across all 1,178 DepMap cell lines.

Surprisingly the cluster defined by the presence of DNA replication initiator factors (enriched in GO:0090329 ‘Regulation of DNA-templated DNA replication’) such as *CDC7*, *DBF4* and *TICRR* (Treslin) also included *CDK8* and *CCNC*. More importantly, when considering all the genes ranked according to their ΔB score, which might correlate with the magnitude of the effect, *CDK8* was the one with the lowest ΔB of -1.377 while *CDK19* and other components of the mediator complex kinase-module, *MED12* and *MED13* were not in the hit list, having a ΔB score ranging from -0.019 to 0.241 (Supplementary Table S1).

These findings very strongly suggest that *CDK8* function might be a major determinant of CDC7i sensitivity and prompted us to test the hypothesis of CDK8 direct involvement in DNA replication.

### CDK8 supports DNA replication in the presence of CDC7 inhibitors

To assess if CDK8 depletion could affect DNA synthesis we generated cell lines conditionally expressing shRNAs directed against CDK8 and, as control, also against its paralog CDK19, which in the context of transcription appear to be mostly redundant (61).

Measuring DNA content and levels of DNA synthesis in individual cells by flowcytometry, we found that neither CDK8 nor CDK19 depletion obviously altered the cell cycle distribution nor EdU incorporation in the cells, whereas treatment with low doses of the CDC7 inhibitor XL413 caused decreased levels of DNA synthesis with cells accumulating in S-phase, in line with previous studies (12, 26) (Fig. 3A-D). The reduction of EdU incorporation upon CDC7i treatment was enhanced in absence of CDK8 while unchanged upon CDK19 depletion (Fig. 3A-D). Consistent with a strong delay in S-phase progression CDK8 depletion together with CDC7 inhibition reduced proliferation (Fig. 3E). Altogether these results indicate that in MCF10A cells CDK8, but not its paralog CDK19, contributes to DNA replication in conditions of limiting CDC7 activity.

**Figure 3:**
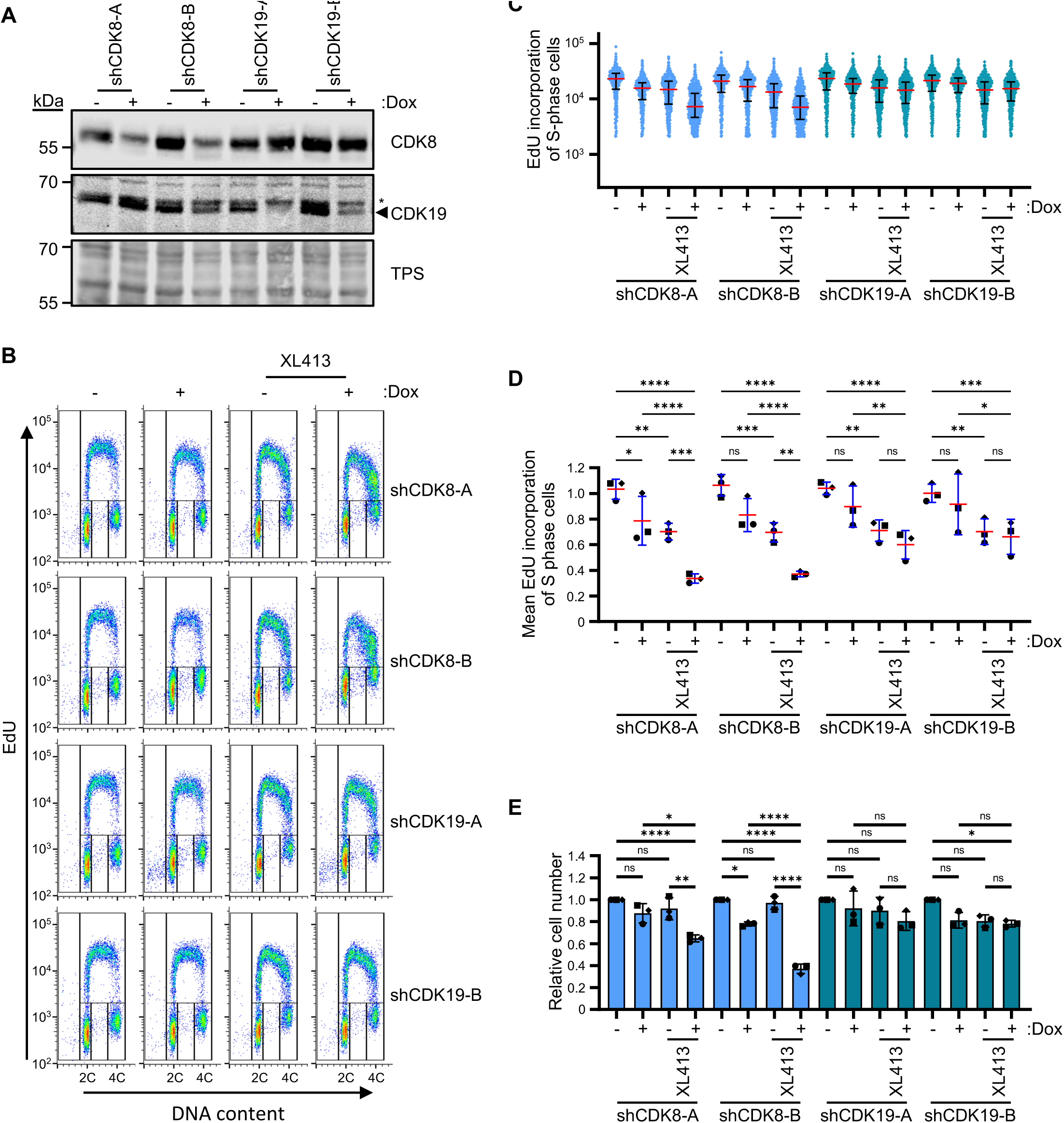
CDK8, but not CDK19, supports replication and proliferation when CDC7 kinase activity is limiting. (A-E) MCF10A cells with doxycycline-inducible expression of shRNA directed against CDK8 or CDK19 were mock treated or treated with doxycycline (Dox, 1 mg/ml) for 48 hours before a further 24 hours in the presence or absence of 1.25 µM XL413. **(A)** Immunoblotting analysis of indicated proteins. TPS (total protein stain) is used as a loading control. Asterisk indicates non-specific band**. (B)** Cells were labelled with EdU for last 30 minutes and DNA synthesis, DNA content and cell cycle distribution were assessed by flow cytometry. **(C)** Fluorescence intensity, proportional to EdU incorporation, in individual S phase cells. Red and black lines indicate the median and the interquartile range extending from the 25^th^ to the 75^th^ percentile. Data is from a single representative experiment. **(D)** Mean fluorescence intensity in all S phase cells from three independent experiments is expressed as a ratio relative to the untreated condition. Red and blue line shows the mean ± SD. Statistical analysis: Two-way ANOVA with Tukey’s multiple comparison post-test; ns, not significant; *P< 0.05; **P< 0.01; ***P< 0.001; ****P< 0.0001. **(E)** Cell number was determined as described in the method details. Mean cell number ± SD (black lines) from three independent experiments is expressed as a ratio relative to the untreated condition. Statistical analysis: Two-way ANOVA with Tukey’s multiple comparison post-test; ns, not significant; *P< 0.05; **P< 0.01; ***P< 0.001; ****P< 0.0001.

To further dissect how CDK8 contributes to DNA replication we moved to a fully pharmacological approach. We first found that adding the CDK8 inhibitor BI-1347 to MCF10A only marginally inhibited proliferation of MCF10A cells, in contrast it strongly reduced cell growth when it was combined with either 1.25 μM or 10 μM of the CDC7i XL413 (Fig 4A-C).

**Figure 4:**
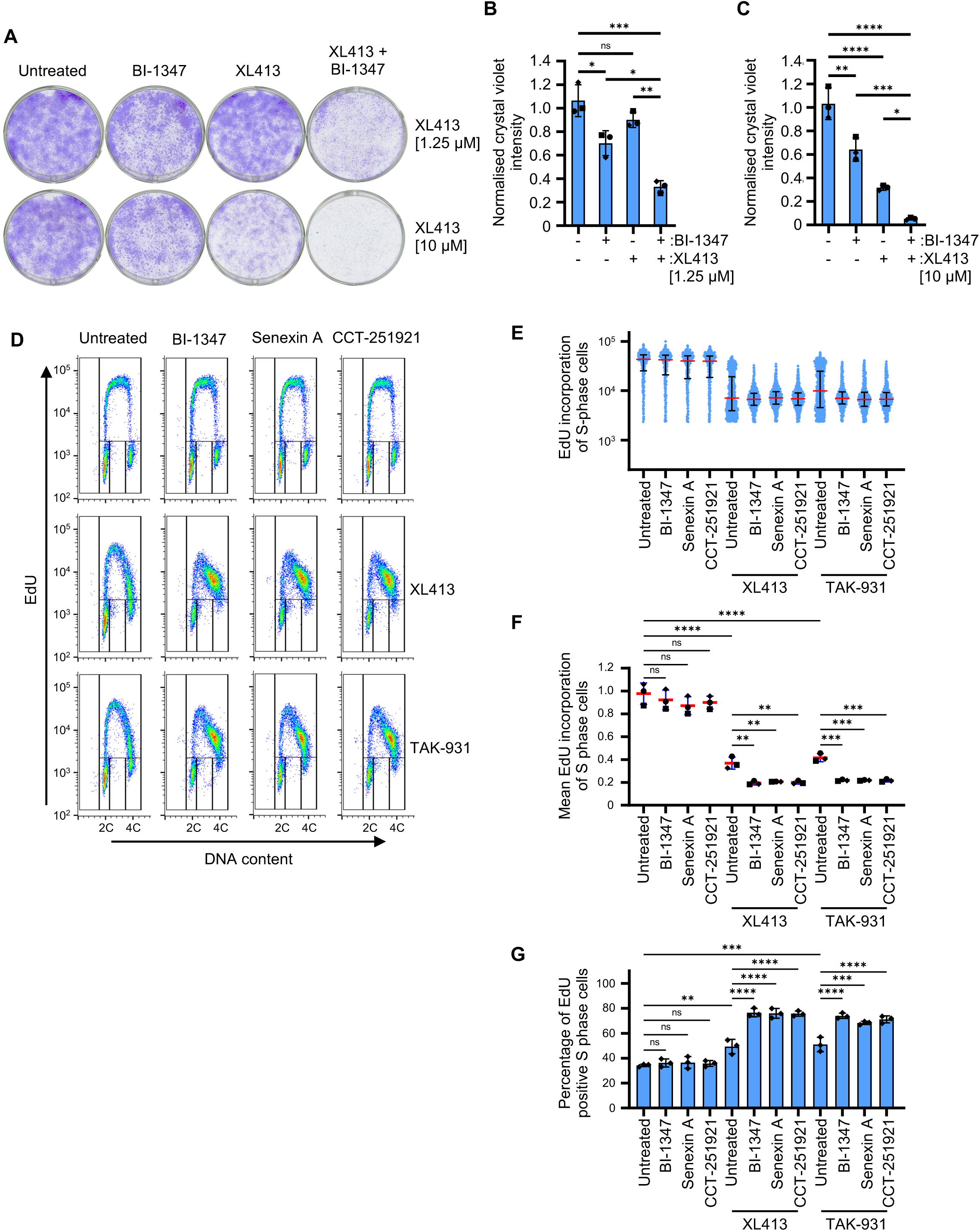
CDK8 and CDC7 inhibitors cooperate to restrict proliferation, DNA replication and S phase progression. (A-C) MCF10A cells were plated at equal low density and either mock treated or treated with 31 nM BI-1347, in the presence or absence of 1.25 µM or 10 µM XL413 over a 7-day period. **(A)** Representative image of plates stained with crystal violet. **(B and C)** Crystal violet intensity was quantified and mean intensity ± SD (black lines) from three independent experiments is expressed as a ratio relative to the untreated condition. Statistical analysis: Two-way ANOVA with Tukey’s multiple comparison post-test; ns, not significant; *P< 0.05; **P< 0.01; ***P< 0.001; ****P< 0.0001. **(D - G)** MCF10A cells were mock treated or treated for 24 hours with three different CDK8 inhibitors: 31 nM BI-1347, 5 µM Senexin A, or 100 nM CCT-251921, alone or in combination with either 10 µM XL413 or 0.3 µM TAK-931 CDC7 inhibitors. **(D)** Cells were labelled with EdU for last 30 minutes and DNA synthesis, DNA content and cell cycle distribution were assessed by flow cytometry. **(E)** Fluorescence intensity, proportional to EdU incorporation, in individual S phase cells. Red and blue lines indicate the median and the interquartile range extending from the 25^th^ to the 75^th^ percentile for each condition. **(D)** Mean fluorescence intensity in all S phase cells from three independent experiments is expressed as a ratio relative to the untreated condition. Red and blue line shows the mean ± SD. Statistical analysis: Two-way ANOVA with Tukey’s multiple comparison post-test; ns, not significant; *P< 0.05; **P< 0.01; ***P< 0.001; ****P< 0.0001. **(G)** The mean percentage (± SD, black lines), of EdU positive S-phase cells for each condition, from three independent experiments is plotted. Statistical analysis: Two-way ANOVA with Tukey’s multiple comparison post-test; ns, not significant; *P< 0.05; **P< 0.01; ***P< 0.001; ****P< 0.0001.

Flow cytometry showed that CDK8 inhibition by BI-1347 treatment for 24 hours did not obviously affect the rate of DNA synthesis or S phase progression unless it was combined with the CDC7 inhibitors XL413 and TAK-931, which resulted in a large fraction of cells accumulating in S-phase (Fig. 4D-G). This was a specific effect of CDK8 inhibition because a variety of chemically distinct CDK8 inhibitors, namely BI-1347, Senexin A and CCT-251921 (Fig. 4D-G), reproduced the same effects of CDK8 depletion with XL413 ruling out potential compound-specific off-target effects.

We then performed similar combination experiments using BI-1347 and XL413 in breast cancer cells MDA-MB-231 and MDA-MB-468, in the osteosarcoma derived U-2 OS cells, and again we observed robust cooperation between the CDC7 and CDK8 inhibitors in suppressing DNA synthesis (Supplemental Fig. S3). These results indicate that CDC7 and CDK8 cooperate to support DNA synthesis. This cooperation is not restricted to one cell-type and might underline a general mechanism for controlling DNA replication in cultured human cells.

### CDK8 promotes origin firing

To gain insight on how CDK8 inhibition restrains DNA synthesis when combined with CDC7 inhibitors, we changed the experimental design in such a way that the effects CDK8i could be assessed with acute rather than chronic inhibitor treatments. Cells were first treated with CDC7i for 16 hours and then we added either CDK8i or DMSO for a further 5 hours. At the end of the experiment the number of S-phase cells was very similar in both situations. However the addition of the CDK8i dramatically reduced EdU incorporation, and this was evident in both early and late S-phase cells (Fig. 5A-C).

**Figure 5:**
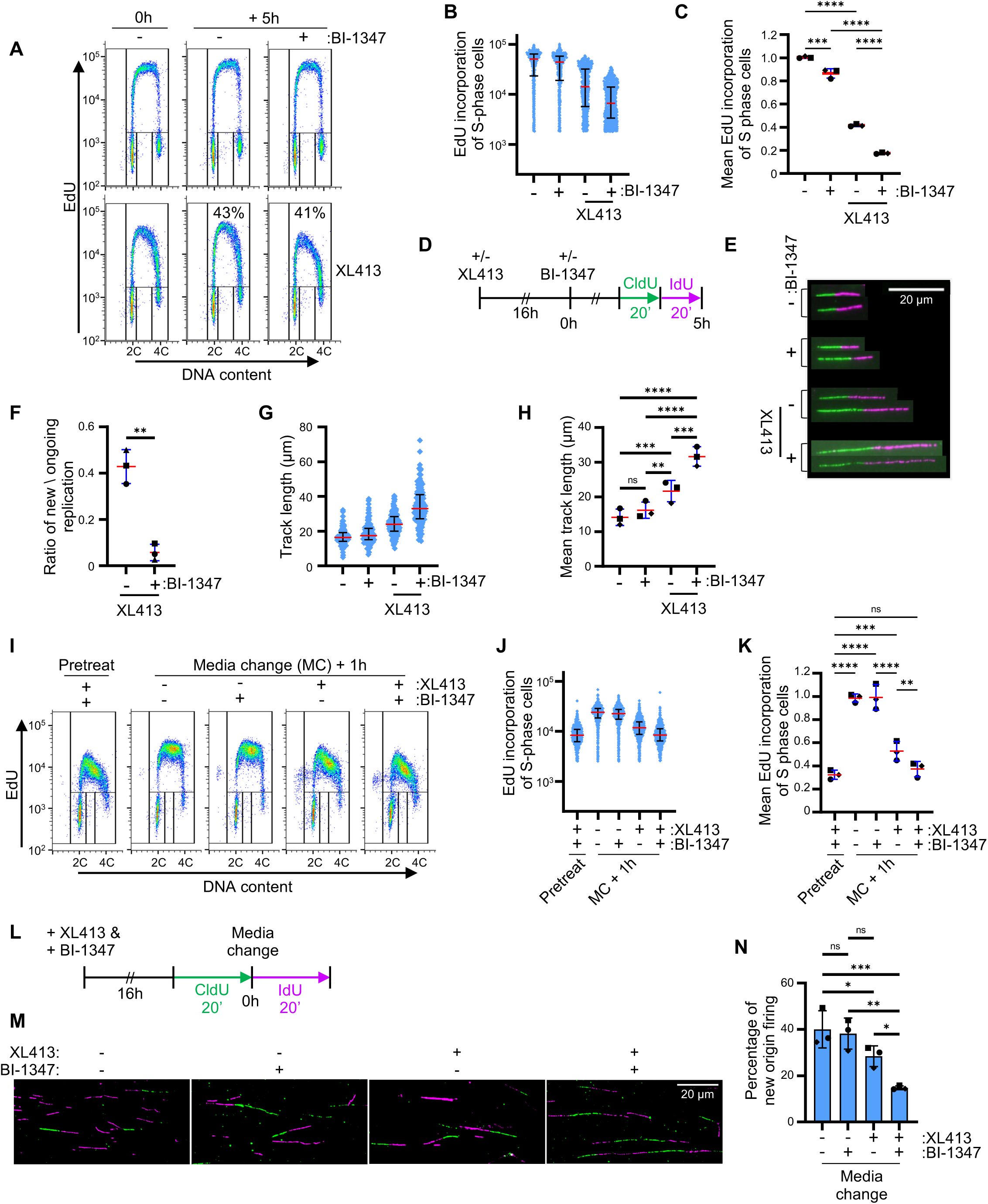
CDK8 stimulates DNA replication origin firing. (A-H) MCF10A cells were either mock treated or treated with 10 μM XL413 for 16 hours before being treated with 31 nM BI-1347 or vehicle for a further 5 hours. **(A)** Cells were labelled with EdU for last 30 minutes and DNA synthesis, DNA content and cell cycle distribution were assessed by flow cytometry. Percentage of EdU positive S phase cells is shown for key conditions. **(B)** Fluorescence intensity, proportional to EdU incorporation, in individual S phase cells. Red and blue lines indicate the median and the interquartile range extending from the 25^th^ to the 75^th^ percentile for each condition. **(C)** Mean fluorescence intensity in all S phase cells from three independent experiments is expressed as a ratio relative to the untreated condition. Red and blue line shows the mean ± SD. Statistical analysis: Two-way ANOVA with Tukey’s multiple comparison post-test; ns, not significant; *P< 0.05; **P< 0.01; ***P< 0.001; ****P< 0.0001. **(D)** Nascent DNA was sequentially labelled with CldU (green) and IdU (magenta) just before cells were harvested for DNA fibre spreads. **(E)** Representative images of DNA fibres for each condition (scale bar, 20 µm). **(F)** ∼200 DNA fibres were scored in each condition. The ratio of new replication (magenta-only tracks) versus ongoing replication (green and magenta tracks) was calculated from three independent experiments and the mean ± SD is plotted (Red and blue lines). Statistical analysis: Unpaired T-test; **P< 0.01. **(G)** The total track length (µm) for at least 200 replication forks is plotted for a single experiment. Red and blue lines indicate the median and the interquartile range extending from the 25^th^ to the 75^th^ percentile for each condition. **(H)** Mean track length ± SD (red and blue lines) from three independent experiments is shown for each condition with statistical analysis: Two-way ANOVA with Tukey’s multiple comparison post-test; ns, not significant; *P< 0.05; **P< 0.01; ***P< 0.001; ****P< 0.0001. **(I-K)** MCF10A cells were first treated with both 31 nM BI-1347 and 10 μM XL413 for 16 hours, then cells were washed and cultured for a further 1h in fresh media or fresh media containing again both inhibitors, BI-1347 only, or XL413 alone. **(I)** Cells were labelled with EdU for last 30 minutes before and after media changes and DNA synthesis and DNA content were assessed by flow cytometry. **(J)** Fluorescence intensity, proportional to EdU incorporation, in individual S phase cells. Red and blue lines indicate the median and the interquartile range extending from the 25^th^ to the 75^th^ percentile for each condition. **(K)** Mean fluorescence intensity in all S phase cells from three independent experiments is expressed as a ratio relative to the untreated condition. Red and blue line shows the mean ± SD. Statistical analysis: Two-way ANOVA with Tukey’s multiple comparison post-test; ns, not significant; *P< 0.05; **P< 0.01; ***P< 0.001; ****P< 0.0001. **(L-N)** MCF10A cells were first treated with both 31 nM BI-1347 and 10 μM XL413 for 16 hours as before, then cells were washed and cultured for a further 20 minutes in fresh media or fresh media containing again both inhibitors, BI-1347 only, or XL413 alone. **(L)** Nascent DNA was labelled with CldU (green), and then with IdU (magenta) for further 20 minutes after media changes. Cells were then harvested and nascent DNA analysed by DNA fibre spreading technique. **(M)** Images of representative DNA fibres for each condition (scale bar, 20 µm). **(N)** The percentage of new origin firing (magenta-only tracks) was scored for ∼200 DNA fibres in each condition in three independent experiments and the mean ± SD is plotted (Black lines). Statistical analysis: Two-way ANOVA with Tukey’s multiple comparison post-test; ns, not significant; *P< 0.05; **P< 0.01; ***P< 0.001; ****P< 0.0001.

The analysis of nascent DNA by fibre microscopy showed that CDK8 inhibition severely reduced the number of new origins fired in the presence of CDC7i. This was accompanied by an increase in the length of newly synthetised replication tracks, indicating that, as expected, replication fork speed increased as a result of an overall lower fork number (Fig. 5D-H).

To corroborate the conclusion that efficient replication requires both CDC7 and CDK8 kinases, we first allowed the cells to accumulate in S-phase by cotreatment with both CDC7 and CDK8 inhibitors and then measured DNA synthesis upon washing off either one or both drugs. DNA synthesis increased immediately after the removal of either CDC7i or CDK8i alone although CDK8i withdrawal was less effective (Fig. 5I-K). DNA fibre analysis showed that the resumption of DNA synthesis resulted from a burst of new origin firing (Fig. 5L-N), directly linking CDK8 activity to origin function.

### CDK8 contribution to origin firing is independent from ATR

We have previously shown that CDC7 inhibition induces a mild replication stress leading to ATR checkpoint pathway activation which further contributes to suppression of origin firing. Under these conditions ATR inhibition allows many origins to be activated, a process that requires CDK1 (26). Thus, we asked if CDK8 inhibition could affect ATR signalling and therefore if its function in origin firing could be related to checkpoint signalling. We observed that CDK8 inhibition alone did not induce FancD2 ubiquitylation nor phosphorylation of Chk1 at serine 345 or ATR at threonine 1989, and kept with almost normal rate of DNA synthesis in these cells (Fig. 6A). However, cotreatment with XL413 increased all replication stress markers measured compared to the treatment with only the CDC7i.

**Figure 6:**
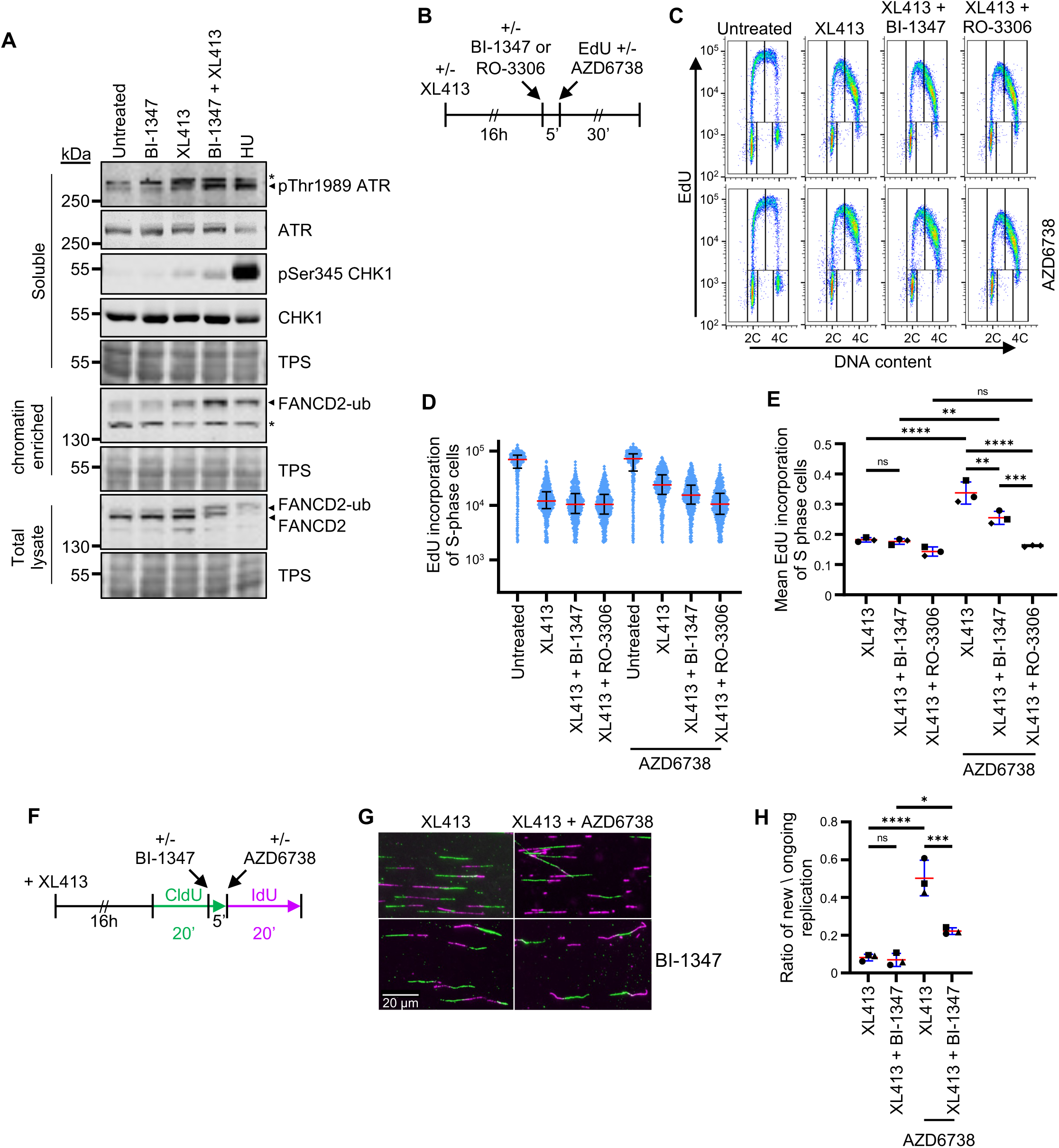
ATR inhibition does not bypass the requirement for CDK8 in origin firing. MCF10A cells were either mock treated, treated with 31 nM BI-1347 alone, with 10 µM XL413 with or in combination of both or with 2mM Hydroxyurea (HU) for 24 hours. **(A)** Whole cell extracts, or soluble and chromatin-enriched fractions were analysed by immunoblotting for the indicated proteins. TPS (total protein stain) is used as a loading control. Asterisks indicate non-specific bands**. (B-E)** MCF10A cells were mock treated or treated with 20 μM XL413 for 16 hours before being further treated with 31nM BI-1347 or with 10 µM of the CDK1 inhibitor RO-3306 for 5 minutes. At this point, EdU was added and half of the cultures received 5 μM of the ATR inhibitor AZD6738 and collected after 30 minutes. **(B)** Outline of experimental design. **(C)** DNA synthesis and DNA content were assessed by flow cytometry. **(D)** Fluorescence intensity, proportional to EdU incorporation, in individual S phase cells. Red and blue lines indicate the median and the interquartile range extending from the 25^th^ to the 75^th^ percentile for each condition. **(E)** Mean fluorescence intensity in all S phase cells from three independent experiments is expressed as a ratio relative to the untreated condition. Red and blue line shows the mean ± SD. Statistical analysis: Two-way ANOVA with Tukey’s multiple comparison post-test; ns, not significant; *P< 0.05; **P< 0.01; ***P< 0.001; ****P< 0.0001. **(F and H)** Cells were treated with 20 μM XL413 for 16 hours before being further treated with 31nM BI-1347 or vehicle for 5 minutes. before the addition of ATR inhibitor AZ6738. Nascent DNA was labelled for 20 minutes with CldU (green) before the addition of ATRi and then for a further 20 minutes with IdU (magenta). **(F)** Outline of experimental design. **(G)** Representative images of DNA fibres in each condition (scale bar, 20 µm). **(H)** The ratio of new origin firing (magenta-only tracks) versus ongoing replication (green and magenta tracks) was calculated from ∼200 DNA fibres for each condition in three independent experiments and the mean ± SD is plotted (Red and blue lines). Statistical analysis: Statistical analysis: Two-way ANOVA with Tukey’s multiple comparison post-test; ns, not significant; *P< 0.05; **P< 0.01; ***P< 0.001; ****P< 0.0001.

These findings suggested that, potentially, the reduction in DNA synthesis and origin firing by CDK8i in presence of CDC7i, could be indirect effect and caused by unexpected problems at forks, thus reinforcing ATR signalling and further limiting origin activation. To test this hypothesis, we first treated the cells with CDC7i for 16 hours and then added the CDK8i BI-1347 or the CDK1i RO-3306 five minutes before the ATR inhibitor AZD6738. The rate of DNA synthesis was then monitored by flow cytometry. As previously described ATRi induced a burst of DNA synthesis in CDC7i treated cells which was fully suppressed by CDK1i (Fig. 6B-E) (26). Interestingly, this ATR-dependent sudden increase in DNA synthesis was also partially suppressed by CDK8i (Fig. 6B-E). In parallel DNA fibre experiments we observed that the burst of origin firing due to ATR inhibition was indeed also reduced by CDK8 inhibition (Fig. 6F-H).

Altogether, these experiments show that CDK8 is still required for efficient origin firing in absence of checkpoint signalling and suggest a direct role for this kinase in origin activation.

### CDK8 function requires MTBP-CDK8 interaction, with CDK8 and CDC7 phosphorylating MCM4

The origin factor MTBP binds to CCNC, and it is likely recruiting active CDK8 kinase to the pre-RC for origin activation. To test this possibility, we examined whether disrupting MTBP-CCNC interaction would affect DNA synthesis in presence of CDC7i. For this purpose, isogenic Hela_Flp-In cells expressing either one of the siRNA resistant transgenes MTBP wild-type or two CDK8 binding mutants, MTBP-7pm (L620D, P622D, L623D, F632A, V633D, L634D, T635A) and 1D (T687D), that carry non-overlapping amino acid exchanges, were used (Supplemental Fig. S4C).

These cells were transfected with MTBP targeting siRNA and treated with doxycycline to induce the expression of the transgenes and the rate of DNA synthesis was assessed in absence or presence of the CDC7i XL413. Parental Hela cells not expressing a transgene were used as a control. MTBP depletion had a severe effect on replication compared with parental Hela cells treated with control siRNA (Fig. 7A). MTBP-WT and the two CDK8 non-binding mutants rescued replication in DMSO treated cells with only minimal differences in BrdU incorporation, as reported previously (52). In contrast to DMSO treatment, partial CDC7 inhibition strongly reduced the levels of BrdU incorporation in cells expressing the MTBP-7pm and 1D mutants but not in cells expressing MTBP-WT (Fig. 7A-C, Supplemental Fig. S4A-B), thus revealing the functional relevance of MTBP-CDK8 interaction in DNA replication when CDC7 activity is limiting.

**Figure 7:**
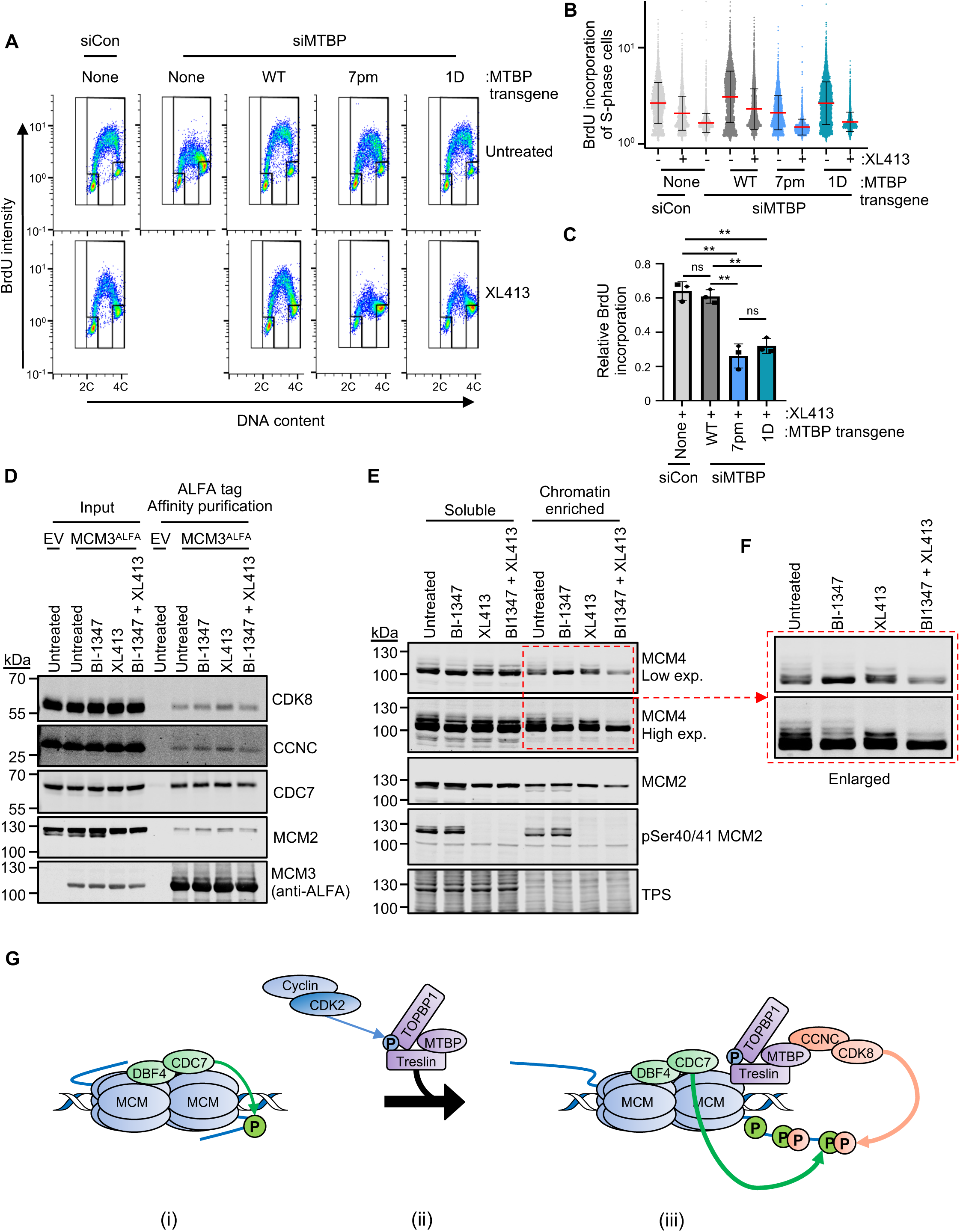
CDK8 acts with MTBP in origin activation and contributes to MCM4 phosphorylation. **(A)** HeLa Flip-In T-Rex cells with doxycycline-inducible expression of siRNA-resistant and C-terminally tagged (3x Flag-TEV2-GFP) MTBP wild type (WT), MTBP-Cdk8 non-binding mutants: seven point mutant (7pm), or single point mutant (1D), or no transgene were transfected with control siRNA (siCon) or siRNA against MTBP (siMTBP) and treated with 1 µg/ml doxycycline for 72 hours. Cells were either mock treated or treated with 2,5 µM XL413 for 24 hours before labeling with BrdU for last 30 minutes. DNA synthesis, DNA content and cell cycle distribution were assessed by flow cytometry. **(B)** Fluorescence intensity, proportional to BrdU incorporation, in individual S phase cells. Red and blue lines indicate the mean ± SD. **(C)** Mean BrdU fluorescence intensity from three independent experiments relative to the control condition. Black lines show the mean ± SD. Statistical analysis: parametric, unpaired, two-tailed student t-test: ns, not significant; **P<0.01. **(D)** HEK 293T/17 cells were transfected with C-terminally ALFA-tagged MCM3 (MCM3^ALFA^), or empty vector (EV). After 24 hours cells were either mock treated or treated with 10 μM XL413 for 16 hours before being treated with or without 31nM BI-1347 for a further 5 hours. ALFA tag affinity purifications were performed on clarified chromatin-enriched cell lysates and analysed by immunoblotting for the indicated proteins. **(E)** HEK 293T/17 cells were either mock treated, treated with 31 nM BI-1347 alone, with 10 µM XL413 alone, or with the combination of both for 24 hours. Soluble and chromatin-enriched fractions were analysed by immunoblotting for the indicated proteins. TPS (total protein stain) is used as a loading control. **(F)** Inset shows and enlargement of the indicated area (red dotted line) of the immunoblot in **(E)**. **(G)** Model for origin activation outlining the contribution of the three kinases. CDC7 directly binds to the double MCM hexamer (DH-MCM) and partially phosphorylates the MCM4 tail. Meanwhile CDK2 promotes the assembly of the TOPBP1-Treslin-MTBP complex which is then recruited to the DH-MCM. CDK8 through its interaction with MTBP is now positioned to phosphorylate MCM4 at S/T-S/T-P sites at the proline proximal S/T. S/T-pS/T-P motives are now favourable substrates for further CDC7 dependent phosphorylation, relieving the inhibitory activity of MCM4 tail and efficient origin firing.

We then transfected HEK 293T cells with a construct expressing MCM3 protein fused to an ALFA-tag. We found that MCM3-ALFA similarly to other MCM subunits, fractionated with chromatin and that in pull down experiments from chromatin-enriched fraction it interacts with MCM2, indicating that MCM3-ALFA could be functionally incorporated within the replicative helicase (Fig. 7D). Notably in these pull downs we also found CCNC, CDK8 and CDC7. As the extracts had been treated extensively with benzonase, it is likely that these findings reflect genuine protein-protein interactions and are not mediated by DNA.

Phosphorylation of MCM4 in its N-terminal tail by CDC7 has long been associated to active initiation and it can be detected by multiple bands with altered mobility in SDS-page (8, 17, 19, 26, 62). We treated HEK 293T cells with either CDK8, CDC7 or both inhibitors and soluble and chromatin associated proteins were separated and analysed by western blotting. Surprisingly we observed that CDK8 inhibition markedly reduced the number of MCM4 phospho-isoforms, even more evidently than CDC7 inhibition and with a different pattern (Fig. 7E-F and Supplemental Fig. S4D-E)). Notably cotreatment with CDC7 and CDK8 inhibitors caused further reduction in MCM4 phospho-dependent mobility shifts. On the other hand, MCM2 phospho-dependent shift and phosphorylation at Ser40/41 were not affected by CDK8 inhibition but solely by CDC7 inhibition.

Altogether our results indicate that both CDC7 and CDK8 kinase are recruited to the pre-replicative complexes to cooperatively phosphorylate MCM4 possibly to activate origins (Fig. 7G).

## DISCUSSION

Despite the process of DNA replication having been investigated for many decades, our understanding of the multiple layers of regulation and how cell respond to its inhibition is still incomplete.

We have embarked in finding the genes that reduce proliferation when origin firing is partially compromised, showing that loss of function of CDK8 or CDK8 pharmacological inhibition sensitizes cells to CDC7 inhibition. We provide evidence that both CDK8 and CDC7 contributes to efficient origin firing and that the kinases cooperate in the origin firing and MCM4 phosphorylation, a key step for origin activation.

During this process, we built a bioinformatic pipeline refining the analysis of CRISPR KO screens, virtually removing problematic sgRNAs (57) from libraries and downstream analysis. Such “cleaning process”, while giving confidence on the final hits, can lead to loss of coverage as several genes may become targeted by an insufficient number of independent sgRNAs. In this study we lost approximately 1000 genes, thus it is still possible that a few genes contributing to the CDC7is activity might have been missed. We have used the criteria of genetic co-essentiality to assign hits into potentially functionally related groups. Such analysis is complementary to classic gene ontology as overlap exists when considering outputs and the broad definition of the processes identified. However, only the co-essentiality analysis spotted the link between CDK8 and initiator factors leading us to hypothesise a direct role of CDK8 in DNA replication. We anticipate that such approach could be useful for extracting biological relevant information from other chemo-genetic screens.

Other networks were identified, including DNA repair and Fanconi Anemia (FA), suggesting that proper resolution of crosslinks and/or efficient fork restart by the FA pathway could be important for dealing with reduced levels of initiation. Genes involved in respiration were revealed and it is likely that defective cellular energetics and release of reactive oxygen species (ROS) from mitochondria may lead to enhanced sensitivity to CDC7is. Lastly, centrosomal proteins are recurrent hits suggesting that centrosome function and/or organization is linked to CDC7 activity; while this is still speculative it has been shown that the CDC7i TAK-931 induces centrosome amplification in HeLa cells (16). Further work will be required to experimentally test these predictions.

In MCF10A cells the cooperation with CDC7 is driven by CDK8 and not by CDK19, suggesting a specific role for CDK8 in DNA replication. It is, however, possible that CDK19 and CDK8 are functionally interchangeable and that their relative abundance in the cell is a key determinant for the replication phenotypes. Multiple lines of evidence then indicate that CDK8 role in replication is not related to transcription: 1) neither MED12 nor MED13 were hits in the CRISPR screen, 2) that a very short period of time of CDK8 inhibition is sufficient to suppress origin firing and that 3) CDK8 stimulation of DNA synthesis is fully dependent on its binding to MTBP and it does not occur when MTBP is depleted.

Protein-protein interactions with MTBP and MCMs position CDK8 at the central stage of the replication initiation machinery. We previously showed that MTBP itself is a CDK8 substrate (53) and phospho-proteomics and multi-omics studies indicated that Treslin as well as MCM3 phosphorylation could be dependent on CDK8 activity (61, 63), suggesting the existence of multiple potentially relevant CDK8 targets in replication initiation. We now provide evidence that MCM4 phospho-isoforms are affected by CDC7 and CDK8 inhibition individually and that both CDK8 and CDC7 kinases cooperate in phosphorylating the MCM4 chromatin associated pool. The MCM4 N-terminal tail, which has been proposed as a key regulatory element for MCM helicase activation and is a recognised CDC7 target (64), contains five S/T-P potential CDK sites, two S-D/E potential CDC7 sites, and six S/T-S/T-P sites. Many of these sites are phosphorylated in vivo (https://www.phosphosite.org/proteinAction.action?id=18710), and several shown to be targeted for dephosphorylation by the RIF1-PP1 phosphatase (19). Biochemical and structural studies have revealed that CDC7 kinase has a strong substrate preference for sites with either an aspartic or glutamic acid at position +1, or for peptides containing a S/T-S/T-P motif, where the negative charge in position +1 is provided by pre-phosphorylation by a proline directed kinase (14, 65, 66). It is tempting to speculate that, while CDK8 could directly phosphorylates MCM4 at S/T-P sites, it may also act as the priming kinase for CDC7-dependent phosphorylation at S/T-S/T-P motifs, thus favouring helicase activation. Such a model would then explain the cooperation between CDK8 and CDC7 in origin activation (Fig. 7G).

At present, we can envisage that CDK8 may either increase the probability of pre-RC’s activation in a stochastic manner across all origins in the genome, or locally at specific subsets of origins, an aspect that deserves investigation. Excessive origin firing, which occurs upon ATR and CHK1 inhibition, causes DNA damage that can be suppressed by limiting initiation events with either CDC7 or CDK2 inhibitors (67, 68). Intriguingly another study showed that CDK8/CCNC contributes to the toxicity of ATR inhibitors (69) and, although this was mainly ascribed to a decrease in R-loops, it is possible that the reduced origin firing in CDK8/CCNC deficient cells is an important factor in determining the sensitivity to ATR inhibition.

This study, by describing CDK8 as the third kinase directly involved in initiation, opens the path to future studies addressing the mechanistic aspects of origin activation by the coordinated action of CDC7, CDK8 and CDK2 and how genome replication is achieved in different cell types and organisms or when their respective kinase activities become deregulated.

Finally, these results have important implication for the development of both CDC7 and CDK8 inhibitors suggesting that a combination therapy for cancers involving CDC7 and CDK8 inhibitors warrants investigation.

## Acknowledgements

MDR, LF and CS thank Dr. Shirley Hanley of the Flow Cytometry core facility in the Technology Services Directorate (University of Galway) for support and assistance in this work, and Yeray Fernández Reyes for critical reading of the manuscript. DB and AP thank Anika Kroesen for excellent technical assistance.

## Disclosures /Conflict of interest statement

M. D. Rainey and C. Santocanale are inventor on EP4275686A1 patent assigned to the University of Galway with title “Combination therapy for cancer comprising a cdc7 inhibitor and a cdk8 inhibitor or a ccnc inhibitor”.

## Funding

This work was supported by Research Ireland grant 23/FFP-A/11683 and by the Research Ireland Centre for Research Training in Genomics Data Science under Grant number 18/CRT/6214. C. J. Ryan was funded by Research Ireland grant number 20/FFP-P/8641. DB and AP were supported by the FOR2800 research unit of the Deutsche Forschungsgemeinschaft (project grant BO3385/4-1).

## Author contributions

1. M. D. Rainey: Conceptualization, Investigation, Visualization, Writing—original draft, Writing—review and editing, Supervision.
2. S. Bernard: Investigation, Methodology, Resources, Visualization, A. Pravi: Investigation, Visualization,
3. L. Farrell: Investigation, Visualization,
4. D. Boos: Conceptualization, Supervision, Writing—review and editing,
5. J. Ryan: Conceptualization, Supervision, Writing—review and editing,
6. Santocanale: Conceptualization, Funding acquisition, Supervision, Writing—original draft, Writing— review and editing.

## METHODS

### Resource Availability

#### Lead Contact

Further information and requests for resources and reagents should be directed to and will be fulfilled by the Lead Contact, Corrado Santocanale.

#### Materials Availability

All reagents generated in this study will be made available on request, but we may require a payment and/or a completed Materials Transfer Agreement if there is potential for commercial application.

## Data and Code Availability

The NGS data for this study will be made available.

The source data for main and supplemental figures will be made available.

The code for ‘library refinement’ and ‘co-essentiality analysis’ will be made available.

### Cell culture

Cell culture was performed in a Class II Bio-safety cabinet and all cell lines were maintained at 37 °C in a humidified atmosphere containing 5% CO_2_. Cell count and viability were determined using a LunaII automated cell counter (Logos Biosystems) and trypan blue exclusion. Culture media and reagents were from Merck unless stated otherwise. MCF10A cell line (human, female origin) was purchased from ATCC and used without further authentication. MCF10A-EditR cells (stably expressing Cas9) were generated as previously described (12). MCF10A cells and derivatives were cultured using DMEM supplemented with 5% horse serum, 25 ng/ml cholera toxin, 10 µg/ml insulin, 20 ng/ml epidermal growth factor (Peprotech), 500 ng/ml hydrocortisone, 50 U/ml penicillin and 50 µg/ml streptomycin. Lenti-X™ 293T cell line (human, female origin) was purchased from TaKaRa and used without further authentication. Routine culture was performed using DMEM +GlutMAX^TM^-I (ThermoFisher Scientific) supplemented with 10% fetal bovine serum, 50 U/ml penicillin and 50 µg/ml streptomycin. MDA-MB-468 (human, female origin) were cultured in RPMI-1640 medium (Merck) supplemented with 2 mM L-glutamine, 10% fetal bovine serum, 50 U/ml penicillin and 50 µg/ml streptomycin. U-2 OS (human, female origin), MDA-MB-231 (human, female origin), HEK 293T/17 (human, female origin) and HeLa Flip-In T-Rex cell lines (70) were cultured in DMEM (Merck) supplemented with 10% fetal bovine serum, 2 mM L-Glutamine, 50 U/ml penicillin and 50 µg/ml streptomycin. Generation of stable HeLa Flip-In T-Rex cell lines expressing MTBP WT, 7pm (L620D, P622D, L623D, F632A, V633D, L634D, T635A) and 1D (T687D) transgenes was performed as previously described (52, 53).

### CRISPR Pooled Lentiviral sgRNA Library

A plasmid based CRISPR pooled lentiviral sgRNA library that can be used in a modular format was purchased from CELLECTA (CELLECTA: CRISPR Human Genome Knockout Library, Module 1, #sgRNA: 52,549, KOHGW-M1-P; Module 2, #sgRNA: 49,461, KOHGW-M2-P; and Module 3, #sgRNA: 48,578, KOHGW-M3-P). Each module covers approximately 6,300 genes and each gene is targeted by 8x sgRNAs (∼50,000 sgRNAs per module), in total targeting >19,000 protein coding genes. Lentiviral production, functional titer estimation and validation in a pooled genome-wide CRISPR/Cas9 screening were performed according to user manual (CRISPR pooled lentiviral sgRNA libraries; www.cellecta.com) with minor modifications as previously described (26).

#### Pooled Genome-wide CRISPR/Cas9 screen

Aliquots of frozen virus that were of known titre and validated in a previous screen (26) were rapidly thawed at 37°C. MCF10A EditR cells (120 x 10^6^ per module) were transduced with virus for individual modules (Modules I, II and III) in the presence of 5 μg/ml polybrene at a Multiplicity of Infection (MoI) of 0.4 achieving ∼600 fold guide representation. After 24 hours, transduced cells were harvested, replated and selected with puromycin (1 μg/ml) for 6 days. Puromycin resistant cells were harvested and a sample of ∼30 x10^6^ cells taken for analysis by next generation sequencing (NGS) at the start of the screen (day 0). The remaining puromycin resistant cells from each module were cultured independently from each other in untreated and 1.25 µM XL413 treated experimental branches. Throughout the screen cell proliferation and viability were determined by cell counting on a LunaII automated cell counter (Logos Biosystems) using trypan blue exclusion. Cells were cultured, harvested, reseeded and retreated every 4 days to maintain ∼600 fold guide representation. Finally at day 20 for the untreated branch and at day 24 for the XL413-treated branch, cells were harvested and a sample of ∼30 x10^6^ cells taken for analysis by NGS.

### Genomic DNA extraction of pooled genome-wide CRISPR/Cas9 screen cell populations

Cell pellets were resuspended in 6 ml of Buffer P1 (50 mM Tris-Cl, pH 8.0, 10 mM EDTA, Qiagen) and lysed by addition of SDS to final concentration of 1 % (v/v). Samples were sonicated at 10 % amplitude for 5 seconds (Branson, digital sonifier) before addition of proteinase K (40 µg/ml, Qiagen) followed by gentle mixing and incubation at 55°C overnight. Samples were cooled to 37°C and following addition of RNaseA (100 µg/ml, Qiagen), incubated for 30min. Samples were chilled on ice before addition of 2 ml of ice cold 7.5 M ammonium acetate (Sigma) followed by two repeat 20 second vortex-pulses and centrifugation at 4,000 g for 10 minutes. The supernatant was transferred to a 15 ml conical tube and genomic DNA was precipitated by addition 6 ml of Isopropanol followed by mixing and centrifugation at 4,000 g for 40 minutes. Genomic DNA was washed with 6 ml of 70 % (v/v) Ethanol, centrifuged at 4,000 g for 10 minutes and the genomic DNA pellet air-dried before being resuspended in 220ul of DNase/RNase free water and quantitated on a Nanodrop.

### sgRNA library amplification

Genomic DNA was subject to a first round PCR using multiple reactions, with a maximum of 50 µg of genomic DNA in each 100 µl PCR reaction, with 0.2 mM of each dNTP, 0.3 µM of each primer (pRSG16-Forward and pRSG16-Reverse: see key resources table), and 1x Titanium *Taq* DNA polymerase in 1x Titanium *Taq* buffer. Genomic DNA was initially denatured at 94°C for 4 minutes, followed by 18 cycles of 94°C for 45 seconds, 63°C for 15 seconds, 72°C for 35 seconds, 1 cycle of 72°C for 4 minutes and hold at 4°C. All 1^st^ round PCR reactions for a sample were pooled, mixed and a 5nl aliquot used directly for 2^nd^ round PCR in a 100 µl final volume, as described before, except using 0.5 µM of each primer designed to add Illumina: flow-cell adaptors; barcodes and annealing sites for sequencing primers (NGS-Fwd and NGS-Rev: see key resources table). The 2^nd^ round PCR was subject to PCR clean-up (Macherey-Nagel), to concentrate the DNA into a 50 µl volume, followed by gel purification (Macherey-Nagel) of a DNA product of ∼293bp from a 2 % (w/v) agarose DNA gel. Gel purified libraries were sent to Novogene for Illumina NGS analysis using a NovaSeq 6000 PE150, paired-end reads and 30Gb of raw data per sample.

### NGS data analysis

FASTQ files obtained by NGS were analysed using MAGeCK version 0.5.9.2 and MAGeCKFlute pipeline version 2.6.0 (58, 71) as outlined below. Cutadapt, a bioinformatics tool to find and remove adapter sequences was used to perform adaptor trimming (72). MAGeCK was then used to map all trimmed sgRNAs and associated sequencing reads to the CELLECTA sgRNA reference library and perform a quality control assessment of the sequencing quality and screen data. Negative control sgRNAs in the CELLECTA library were used by MAGeCK to normalize the read counts between samples. Percentage sgRNA and gene coverage for the *in vitro* screen are reported in Supplemental Fig.1D. For gene hit identification, *in silico* refining of the sgRNA library was performed to eliminate ‘poor quality sgRNAs (defined below) to reduce the possibility of calling false hits. The pipeline to refine the sgRNA library was developed in R version 4.3.2 and using Bioconductor version 3.18 (73, 74). The sgRNA alignment and annotation pipeline was partly adapted from DeKegel & Ryan (75). Human Reference Genome 38 (GRCh38) was indexed using RBowtie version 1.42.0. All sgRNA sequences were aligned using crisprBowtie version 1.6.0 to the GRCh38 with a maximum of 2 mismatches allowed (76, 77). Poor quality sgRNA sequences were defined as: (1) sgRNA that aligned perfectly to multiple locations in the genome (multi-targeting sgRNAs); and (2) sgRNA that aligned with single or double mismatches at the terminal two base positions located distal to the Protospacer Adjacent Motif (PAM) site, and associated with off-target activity (previously associated with off-target activity (57). The specific criteria to define PAM-distal mismatch sgRNA in this study is when the single or double mismatches occur at positions 1 and 2 in the Watson (+) strand and positions 19 and 20 in the Crick (-) strand. All sgRNAs with potential multi-targeting and off-targeting activity were removed from the Cellecta library prior to analysis with MAGeCK. The Consensus Coding Sequence (CCDS) project, a dataset containing all identically annotated protein-coding regions in human and mouse reference genomes, was used to annotate all sgRNAs based on their exon location in the target gene (78). The negative control and intron control sgRNAs were not annotated and were assessed only for multi-targeting and off-targeting activity. This assigning of sgRNAs to exons was performed using GenomicRanges version 1.54.1 (79). This annotation was used to remove all sgRNAs that do not target the exon of a protein-coding gene. The gene symbol corresponding to each sgRNA ID was updated based on the actual mapping between the sgRNA and the target gene. The Human Gene Nomenclature Committee (HGNC) dataset was used to select all sgRNAs targeting only protein-coding genes (80). To ensure sufficient representation of sgRNA number per gene, all genes with fewer than three sgRNAs (1-2 sgRNAs) were removed from the Cellecta library prior to further processing. The sgRNA library refining process yielded a virtual sub-library containing only on-target sgRNAs with no predicted off-target activity. The refined sgRNA library was then used to obtain the sgRNA abundance read count using MAGeCK ‘*count’* command.

Subsequently, sgRNA read count data from the day 0 vs day 20 untreated and day 0 vs day 24 XL413-treated (1.25 µM) conditions were modelled using a negative-binomial distribution by MAGeCK-MLE to calculate a beta score for each targeted gene. Beta score normalization was performed by MAGeCK-Flute, using a previously published list of core essential genes (58), to accommodate the effect of the CDC7 inhibitor on decreased cell proliferation throughout the screen. No copy number correction was applied as the MCF10A EditR cell line used in the screen are near diploid. The normalized control beta score was subtracted from normalized treatment beta score, for each gene target, and the genes were ranked and plotted based on this differential beta score. Gene hits were identified by using the ‘*CutoffCalling’* function in MAGeCKFlute with +/- 3x standard deviation of the mean differential beta score were applied for each of the individual modules. Gene hits identified across the 3 individual modules were collated as genes that upon loss modulate cell proliferation to CDC7 inhibition and were further analysed as a cohort.

### Gene Ontology

Gene Ontology analysis was performed using WebGestalt (WEB-based GEne SeT AnaLysis Toolkit) by submitting a list of gene symbols for the hits alongside a reference set of gene symbols targeted by the CRISPR screen (∼18,029 genes) and adopting the following criteria: Over-representation analysis of *Homo sapiens* genes was performed against the Biological Process (noRedundant) gene ontology database using k-Medoid redundancy removal to report ∼15 clusters. A multiple test adjustment was applied by using Bonferroni correction with significance level ≤ 0.05. Gene Ontology identifiers and terms are reported or summarised as multiple variable plots (GraphPad Prism 10.2.1) alongside data for fold enrichment, the number of genes contributing to a GO term and the adjusted p-value (p-adjust).

### Co-essential genetic interaction network

The co-essential genetic interaction network between CRISPR-KO hits was established by using the Cancer Dependency Map (DepMap) 24Q4 CRISPR-KO dataset. DepMap is a resource of CRISPR-KO and RNAi genetic perturbations across thousands of cancer cell lines spanning multiple cancer subtypes (81). The DepMap 24Q4 CRISPR-KO gene effect score (CRISPRGeneEffect.csv) and corresponding cell lines metadata (ScreenSequenceMap.csv) were retrieved from DepMap portal. The DepMap data processing and analysis was performed in Python version 3.13.0. The gene effect score from DepMap 24Q4 were filtered using Pandas version 2.2.3 (82). The filtering criteria were as follows: only cell lines using the Avana (Broad Dependency Map) library were selected, and only genes with a non-missing gene effect score in all cell lines were obtained. This filtering process yields a gene effect score matrix from 1,178 cell lines and 17,916 protein-coding genes. The matrix was analysed for co-essentiality using the Generalised Least Squares (GLS) method as described in Wainberg et al. (59). The GLS was performed using an adapted python script (*gene_pairs.py*) from Wainberg et al. (2021) and Python packages Numpy version 2.1.3 and Scipy version 1.14.1 (59, 83). The GLS resulted in a 17,090 x 17,090 matrix of GLS p-values and sign of correlation. The GLS p-values and correlation matrix were stacked using Pandas ‘*stack’* function to obtain all gene pairs. False Discovery Rate (FDR) was estimated from GLS p-values using the Benjamini-Hochberg multiple-testing correction implemented in Statsmodels version 0.14.4 (84, 85). Co-essential gene pair relationships between hits from the CDC7i CRISPR-KO screen were identified by selecting pairs with significant co-essential relationships (p<0.05 with FDR < 10%) where both genes are hits from the CDC7i CRISPR-KO screen, and the direction of GLS correlation is positive (+1). This co-essential genetic interaction network was then visualized in Cytoscape version 3.10.2, software to visualize biomolecular interaction networks (86). To identify functionally related subnetworks within this larger network, we manually identified connected components with 3 or more subunits (18 subnetworks). The largest single connected component was further broken into two distinct subnetworks because, as evident in Figure 2, it consisted of two largely non-interacting gene sets (one causing resistance, the other causing sensitivity). This resulted in a total of 19 subnetworks, which were annotated using Gene Ontology enrichment analysis that was performed in WebGestalt and using the same settings as described in the previous section. The network layout in Cytoscape was optimized for visual clarity using yFiles layout algorithms version 1.1.5(87). The MAGeCK beta scores of XL413 treated versus untreated were plotted using Tidyverse 2.0.0.

### Cell proliferation assay

MCF10A cells were seeded at 4,000 cells per well of a 6 well plate and after 24 hours were treated with inhibitors before being incubated for 7 days at 37 °C with 5% CO_2_. Media was removed from the cells which were then washed with PBS and air dried before being fixed with 100 % methanol for 1 hour. Following removal of methanol the plates were air-dried, stained with crystal violet staining solution (0.2% crystal violet, 25% methanol) for 1 hour at room temperature, washed with ddH_2_O and air-dried. Plates were scanned using a HP Scanjet 4050, to generate images of crystal violet stained cells, and crystal violet staining intensity was acquired using the Odyssey infrared imaging system and Image Studio software (LI-COR Biosciences).

### Flow cytometry

To analyse the cell cycle and DNA synthesis, nascent DNA was labelled by incubating cells with 10 µM EdU for 30 minutes prior to harvest. Cells were centrifuged at 450 x g for 5 minutes, washed with PBS and resuspended in PBS containing 70% (v/v) EtOH for 1 hour at -20 °C. Cells were centrifuged and washed with PBS before incorporated EdU was labelled by incubating cells in click reaction buffer (PBS containing 10 µM 6-Carboxyfluorescine-TEG-azide, 10 mM Sodium-L-ascorbate, 2 mM Copper-II-Sulphate) for 30 minutes protected from light. Cells were centrifuged and washed with PBS containing 1% BSA, 5% Tween 20 and then DNA stained in PBS containing 1% BSA and 1 µg/ml DAPI. Fluorescence intensity data for 6-Carboxyfluorescine (488_530_30 nm) and DAPI (405_450_50 nm) were acquired for 10,000 single cells on a BD FACS Canto II and analysed using FlowJo software v10. BrdU pulse-labelling and staining using anti-BrdU-FITC and propidium iodide (PI) were performed as previously described (52). Flow cytometry was performed on a MACSQuant flow cytometer (Miltenyi Biotec), and data were analyzed using FlowJo v10. For quantification of relative mean BrdU incorporation, the mean BrdU intensity of S-phase cells was background-subtracted using the intensities of G1 and G2/M cells. The resulting value was then normalized to the mean BrdU intensity of the corresponding DMSO control. Data visualization and statistical analysis were performed using GraphPad Prism 8.

To further analyse fluorescence intensity, proportional to EdU incorporation, for individual cells, gates were applied on biparametric dot plots (DAPI-EdU) to select for EdU positive S phase cells using FlowJo v10. Fluorescence intensity (488_530_30) values per cell were exported and plotted as scatter dot plots using GraphPad prism. For comparisons of mean fluorescence intensity of a cell population across three biological repeat experiments, all fluorescence intensity values were normalised to the minimum and maximum EdU fluorescence intensity of the untreated control sample of each respective experiment to account for absolute differences in EdU staining. The mean fluorescence intensity values of these cell populations across three biological repeat experiments are expressed as a ratio relative to a control sample and plotted as a scatter dot plot using GraphPad prism.

### Constructs

The pRSGTEP-U6Tet-sg-EF1-TetRep-2A-Puro plasmid (SVCRU6TEP-L, Cellecta) was purchased as a linear plasmid for cloning, lentiviral delivery and inducible expression of sgRNAs under the control of the tetracycline-regulated mammalian expression system. The partially complementary oligonucleotides: StufferA-fwd (5’-ACCGGAGTCTTCTTTTTTGAAGACAC-3’) and stufferA-rev (5’-CGAAGTGTCTTCAAAAAAGAAGACTC-3’) were annealed to generate double-stranded DNA with single-stranded DNA overhangs complementary for cloning. Circularisation of pRSGTEP-U6Tet-sg-EF1-TetRep-2A-Puro-StufferA generated two asymmetric BbsI restriction sites that allow for plasmid propagation, linearization and cloning of duplexed oligonucleotides capable of inducible expression of either shRNAs or sgRNAs. Oligonucleotides coding for shRNAs (supplementary reagents table) were annealed and cloned into the BbsI restriction sites of the modified pRSGTEP-U6Tet-sg-EF1-TetRep-2A-Puro-stufferA plasmid to generate pRSGTEP-shRNA transfer plasmids: pRSGTEP-shCDK8-[A], pRSGTEP-shCDK8-[B], pRSGTEP-shCDK19-[A], pRSGTEP-shCDK19-[B] plasmids.

All constructs were propagated in NEB stable *E. coli* (C3040I, NEB). The pcDNA3.1-MCM3^ALFA^ plasmid was synthesised as a FLASH gene (GenScript). The start codon for the MCM3 coding sequence (Accession number: NM_002388) was preceded by a Kozak sequence (gctagcgccgccacc) and the DNA sequence (ggttccccgagccgcctggaagaagaactgcgccgccgcctgaccgaaccg) was inserted immediately upstream of the MCM3 stop codon and encodes for an in-frame linker and ALFA tag (amino acid sequence: GSPSRLEEELRRRLTEP). The FLASH gene insert was flanked by NheI and EcoRI restriction sites at the 5’ and 3’ ends respectively.

### Small scale lentiviral packaging

Lenti-X 293T cells were seeded at 140,000 cells per well in a 6-well plate. After 24 h transfections were performed using a 1:2 ratio w/v of DNA to polyethyleneimine “MAX” MW 40,000 (1 mg/ml, Polysciences). Lentiviral Packaging Plasmids (2 μg; Cellecta) and pRSGTEP-shRNA transfer plasmids (0.4 μg of: pRSGTEP-shCDK8-[A]; pRSGTEP-shCDK8-[B]; pRSGTEP-shCDK19-[A]; or pRSGTEP-shCDK19-[B]) were added to 100 μl of 150 mM NaCl prior to mixing with polyethyleneimine (4.8 μl) that was diluted separately in 100 μl of 150 mM NaCl. Reagents were vortexed and incubated at room temperature for 20 min before dropwise addition to cells. Media was changed after 24 h and transfected Lenti-X 293T cells were allowed to produce virus for a further 24 h. Media (2 ml) was harvested from cells, centrifuged (1,500 rpm for 5 min) and passed through a 0.45 μm sterile filter disc before aliquots were stored at -80°C.

### Generating MCF10A cell lines with inducible shRNA expression

MCF10A EditR cells were plated at 120,000 cells per well and after 24 h were transduced with viral particles (pRSGTEP-shCDK8-[A], pRSGTEP-shCDK8-[B], pRSGTEP-shCDK19-[A] or pRSGTEP-shCDK19-[B]) at a Multiplicity of infection (MoI) of 0.1-0.2. Media was changed after 24 h and a polyclonal population of transduced cells were selected using 0.7 µg/ml puromycin (ant-pr-1, InvivoGen) and included: MCF10A-shCDK8[A], MCF10A-shCDK8[B], MCF10A-shCDK19[A], MCF10A-shCDK19[B]. Expression of shRNA was routinely induced by treating cells with 1mg/ml Doxycycline (D3447, Merck) over a period of 48 to 72 hours.

### Transient transfections

HEK 293T/17 cells were seeded at 5×10^6^ cells per flask and after 24 h transfections were performed using a 1:3 ratio w/v of DNA to polyethyleneimine. pcDNA3.1-MCM3^ALFA^ plasmid (20 μg) was added to 100 μl of 150 mM NaCl prior to mixing with polyethyleneimine (60 μl) that was diluted separately in 100 μl of 150 mM NaCl. Reagents were vortexed and incubated at room temperature for 20 min before being mixed with media and added to cells. Media was changed after 24 h and cells were treated as required.

### siRNA transfections

siRNA-mediated depletion using siRNAs targeting MTBP or a control siRNA (GL2) were performed as previously described (52). Transfections were performed according to the manufacturer’s instructions.

### Protein Preparations

Cells were harvested and fractionated by resuspension in TGN buffer (50 mM Tris-HCl (pH 7.5), 150 mM NaCl, 50 mM Sodium β-glycerophosphate, 0.5% (v/v) Tween-20, 0.2% (v/v) Nonidet P-40, 2 mM NEM, 1 mM Sodium Orthovanadate, 10 mM NaF, 1:100 (v/v) protease and phosphatase inhibitors (Fisher Scientific) for 20 minutes on ice, before being centrifuged at 10,000 x g for 5 minutes at 4°C. The supernatant was transferred to a microfuge tube (Soluble) and the pellet was washed with TGN buffer and centrifuged as before. The pellet was resuspended in TGN buffer supplemented with 1 mM MgCl_2_ and benzonase (125 U/ml) and incubated for 30 minutes on ice before sonication for six cycles (10 seconds on, 30 seconds off) on high setting (BioruptorTM, Diagenode) to further fragment DNA (chromatin-enriched fraction). For ALFA tag affinity purifications the chromatin-enriched fraction was clarified (clarified chromatin-enriched fraction) by centrifugation at 16,000 x g for 5 minutes at 4°C. Protein content was determined using the Bio-Rad DC™ Protein Assay Kit II (5000112, BioRad) and 20 µg of extract was denatured at 95°C in 1x Laemmli buffer and analyzed by SDS-PAGE and immunoblotting. For total cell lysates, cell pellets were resuspended in 1x Laemmli buffer and sonicated, as described above, to fragment DNA before being denatured at 95°C and analyzed by SDS-PAGE and immunoblotting.

### ALFA tag affinity purifications

Clarified chromatin-enriched fractions, obtained from TGN cell lysates of samples, were precleared on agarose resin for 1 hour with rotation (20 rpm) at 4°C, followed by centrifugation at 1,000 x g for 5 minutes. Matched amounts of input protein (∼1 to 2 mg) from precleared samples were then incubated with ALFA Selector CE affinity resin (25 µl bed volume, NanoTag Biotechnologies) for 1 hour with rotation at 4°C followed by centrifugation at 1,000 x g for 1 minute. ALFA Selector resin was washed five time with TGN buffer before affinity purified proteins were eluted at 95°C in 1x Laemmli buffer and analyzed by SDS-PAGE.

### Immunoblotting

Proteins were transferred onto nitrocellulose membranes using a wet transfer system. Proteins on membranes were stained with fast green (0.0001% w/v in 0.1% acetic acid) as a total protein stain (TPS), and after mild destaining in distilled water, with gentle rocking for 5 minutes, were analyzed on the Odyssey infrared imaging system at 700 nm (Li-COR Biosciences). Membranes were completely destained in 0.1 M NaOH in 30% Methanol with gentle rocking for 10 minutes before being blocked in 3% skim milk (Sigma) in TBST (10 mM Tris-HCl pH7.5, 150 mM NaCl, 0.05% Tween 20) for 1 h at room temperature. Immunoblotting was performed by primary antibody incubations in blocking buffer overnight at 4°C, followed by three washes in TBST, secondary antibody incubations in blocking buffer for 1 h and then three washes in TBST all at room temperature. Signals were acquired using the Odyssey infrared imaging system and analyzed using Image Studio software. Primary antibodies were diluted in 3% skim milk/TBST: ATR (sc-515173; Santa Cruz Biotech; 1:1000), FANCD2 (ab5360; abcam; 1:1000), CHK1 (sc8408; Santa Cruz Biotech; 1:1000); CDC7 (K0070-3, MBL, 1:1000); and MCM2 (in house: 1:3000), or 1% BSA/TBST: CDK8 (17395, CST, 1:1000); CDK19 (Cdk11 (8B6) sc-517026, SCBT, 1:500); CCNC (68179, CST, 1:1000); MCM4 (ab4459, Abcam, 1:1000); pT1989 ATR (GTX128145, GeneTex, 1:1000); pS345 CHK1 (2348, CST, 1:1000); Anti-ALFA antibody (54963, CST, 1:2000) and pS40/S41 MCM2 (in house, 1:3000). IRDye secondary antibodies (Li-COR): 800CW goat anti-rabbit (926-32211, Li-COR Biosciences, 1:10,000); and 800CW goat anti-mouse (926-32210, Li-COR Biosciences, 1:10,000) were diluted in 3% skim milk/TBST.

### DNA fibre analysis

Cells were treated with small molecule inhibitors before being sequentially pulse-labelled with the halogenated nucleotides, CldU (1^st^ pulse, 50µM) and IdU (2^nd^ pulse, 200 µM), according to the treatment schemes associated with the relevant figures. DNA replication was blocked by addition of 1 mM Thymidine for 2 min prior to harvest. To prepare and analyse DNA fibre spreads cells were resuspended in PBS at 2.5 × 10^5^ cells per ml before 2.5 μl of cell suspension was mixed with 7.5 μl of lysis buffer (200 mM Tris–HCl, pH 7.5, 50 mM EDTA, and 0.5% (w/v) SDS) on a glass slide. After 8 min, the slides were tilted at ∼15°, and the resulting DNA spreads were air-dried and fixed in 3:1 methanol/acetic acid overnight at 4°C. The fibres were denatured with 2.5 M HCl for 1 h, washed with PBS and blocked with 0.2% Tween-20 in 1% BSA/PBS for 30 minutes. The newly replicated CldU and IdU tracks were labelled (for 1.5 h in the dark, at RT) with anti-BrdU rat monoclonal antibody antibodies recognizing CldU (1:100, ab6326 (BU1/75); Abcam) and anti-BrdU IgG1 mouse monoclonal antibody (1:100, 347580 (B44); BD Biosciences) recognizing IdU. Following three PBS washes the fibres were incubated for 1 hour at room temperature in the dark with secondary antibodies: chicken anti-rat Alexa Fluor 488 (1:300, A21470; ThermoFisher), goat anti-mouse IgG1 Alexa Fluor 546 (1:300, A21123; ThermoFisher). After a final PBS wash coverslips were mounted using SlowFade Gold (S36936, Thermofisher) and images were captured with an IX81-Olympus microscope and 60 X oil-immersion objective (UPlanSApo, infinite/0.17/NF26.5). Images were processed using ImageJ software and analysed using the DNAstranding application (88).

### Quantification and Statistical analysis

GraphPad PRISM 10.2.1 for windows was used to generate all graphs and perform statistical test. Information about the statistical tests and the number of repeat experiments analysed are provided in the figure legends.

## Reagent table

### KEY RESOURCES TABLE

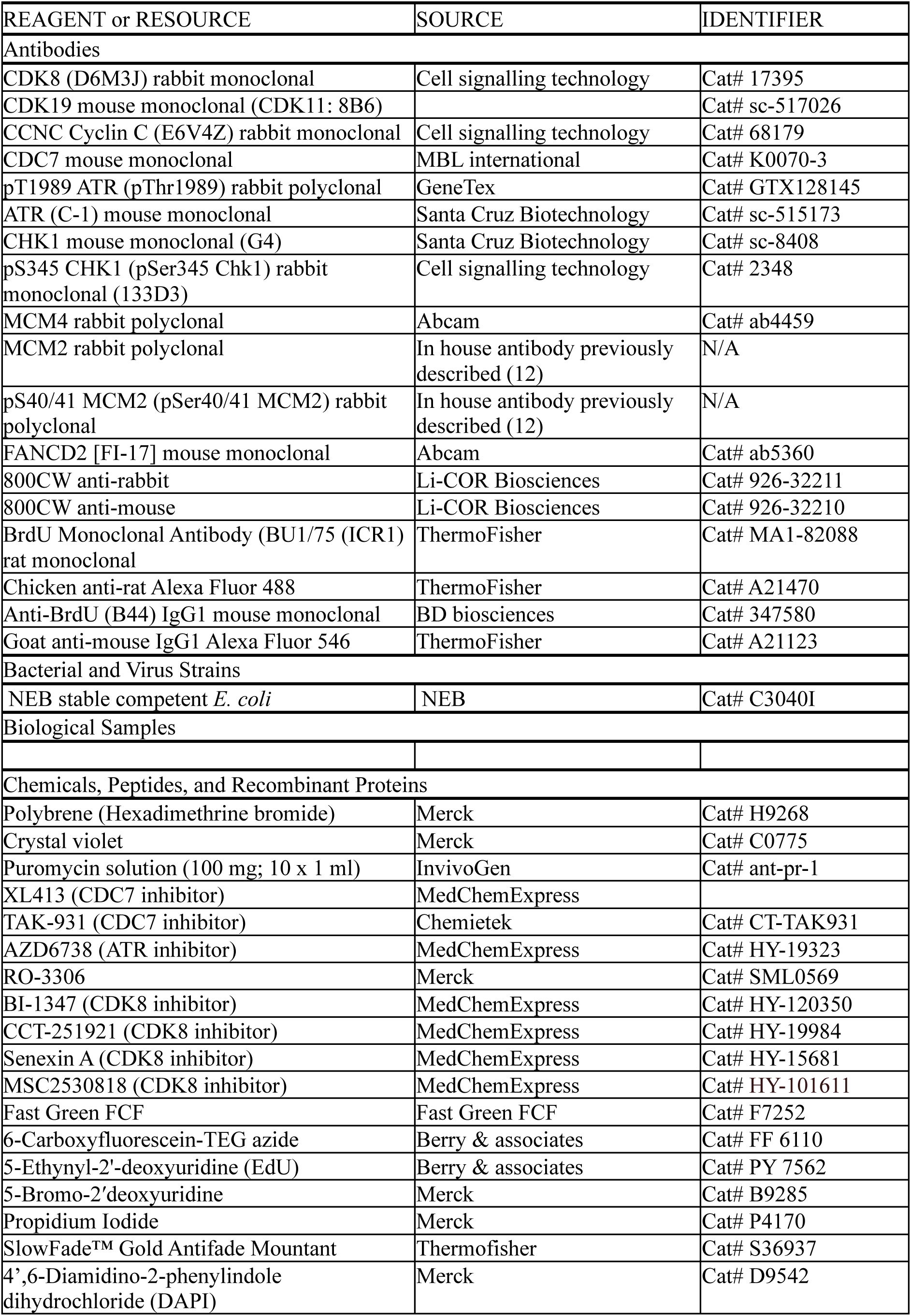
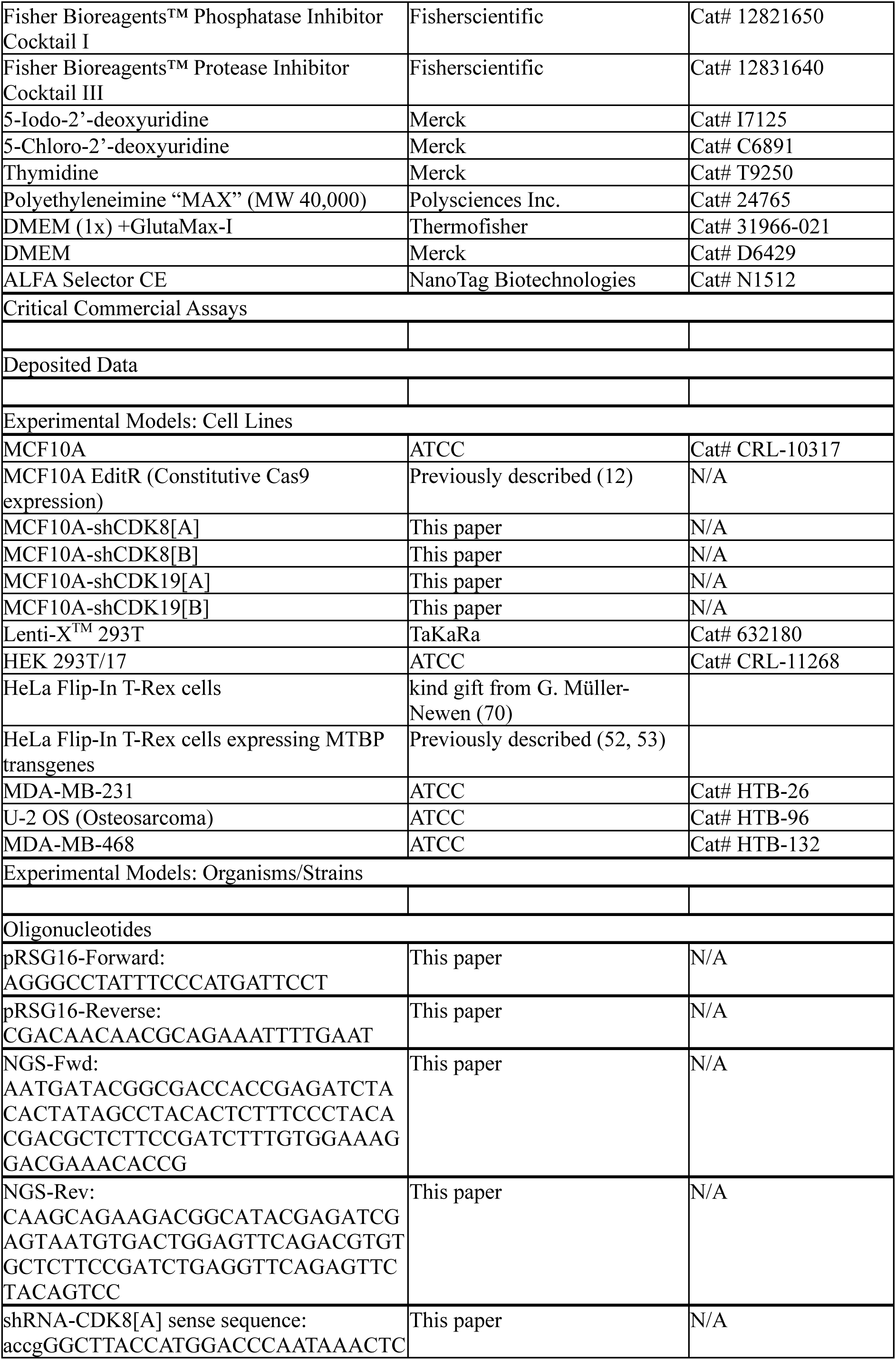
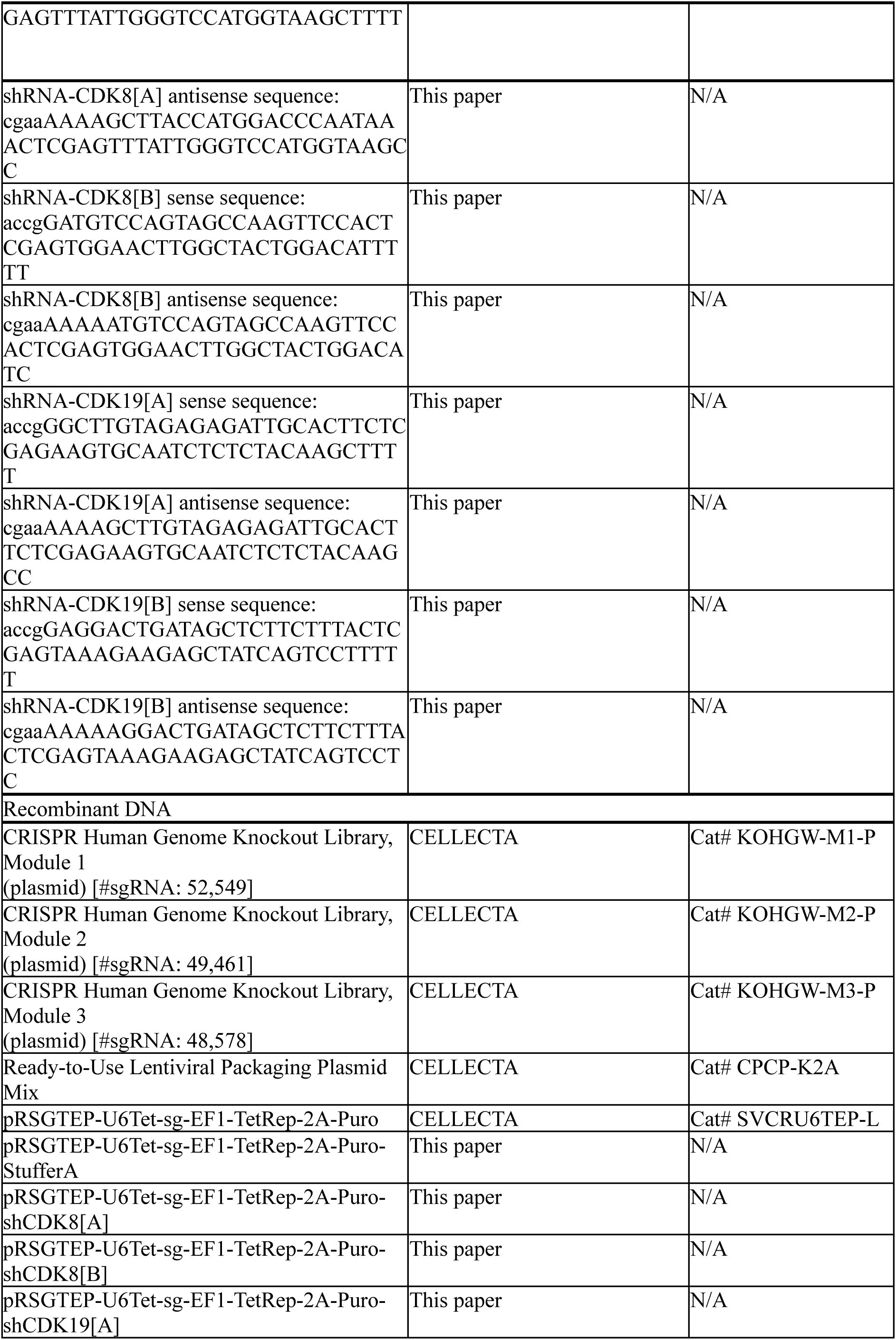
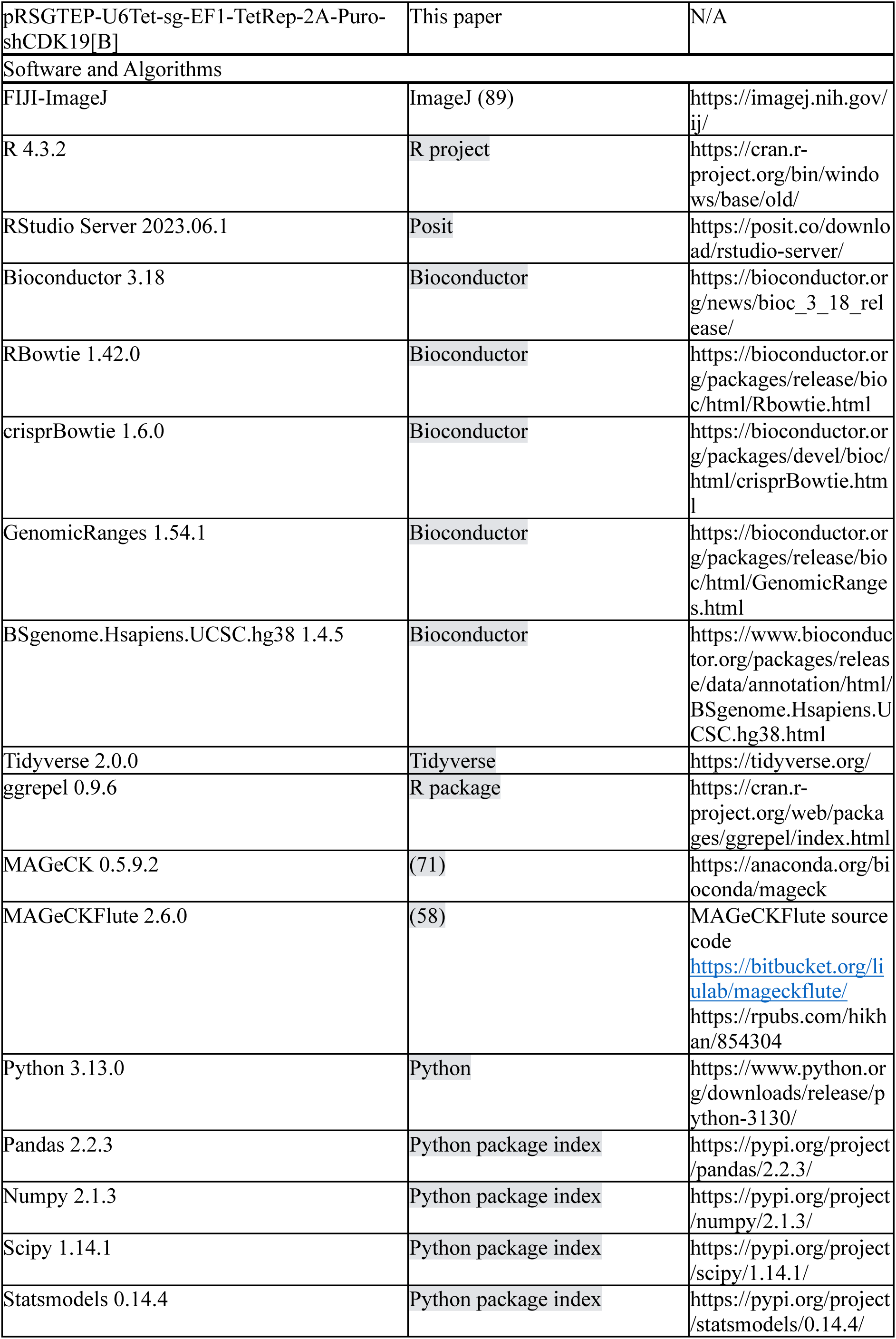
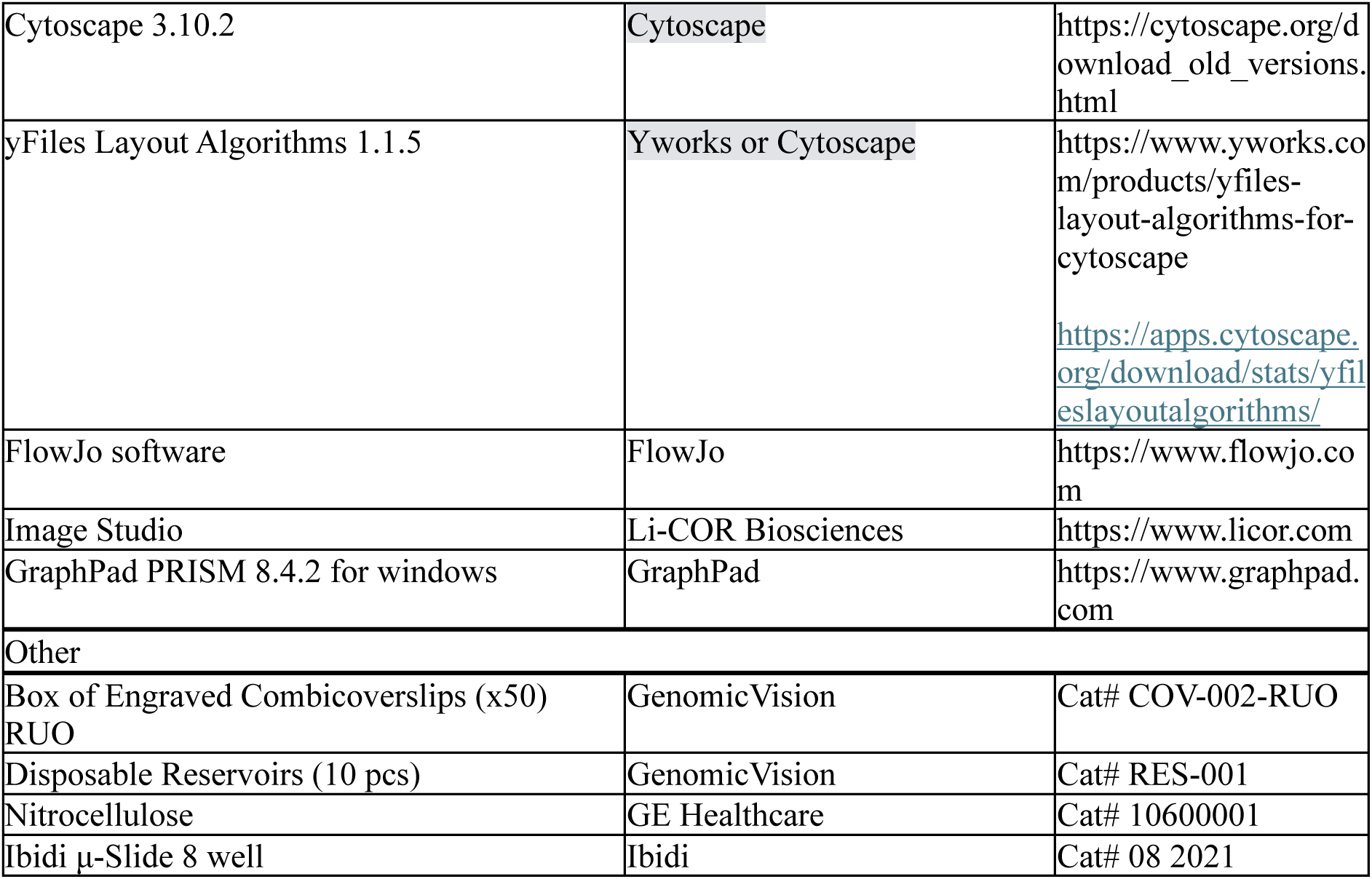

## SUPPLEMENTAL FIGURE LEGENDS

**Supplemental Figure 1:**
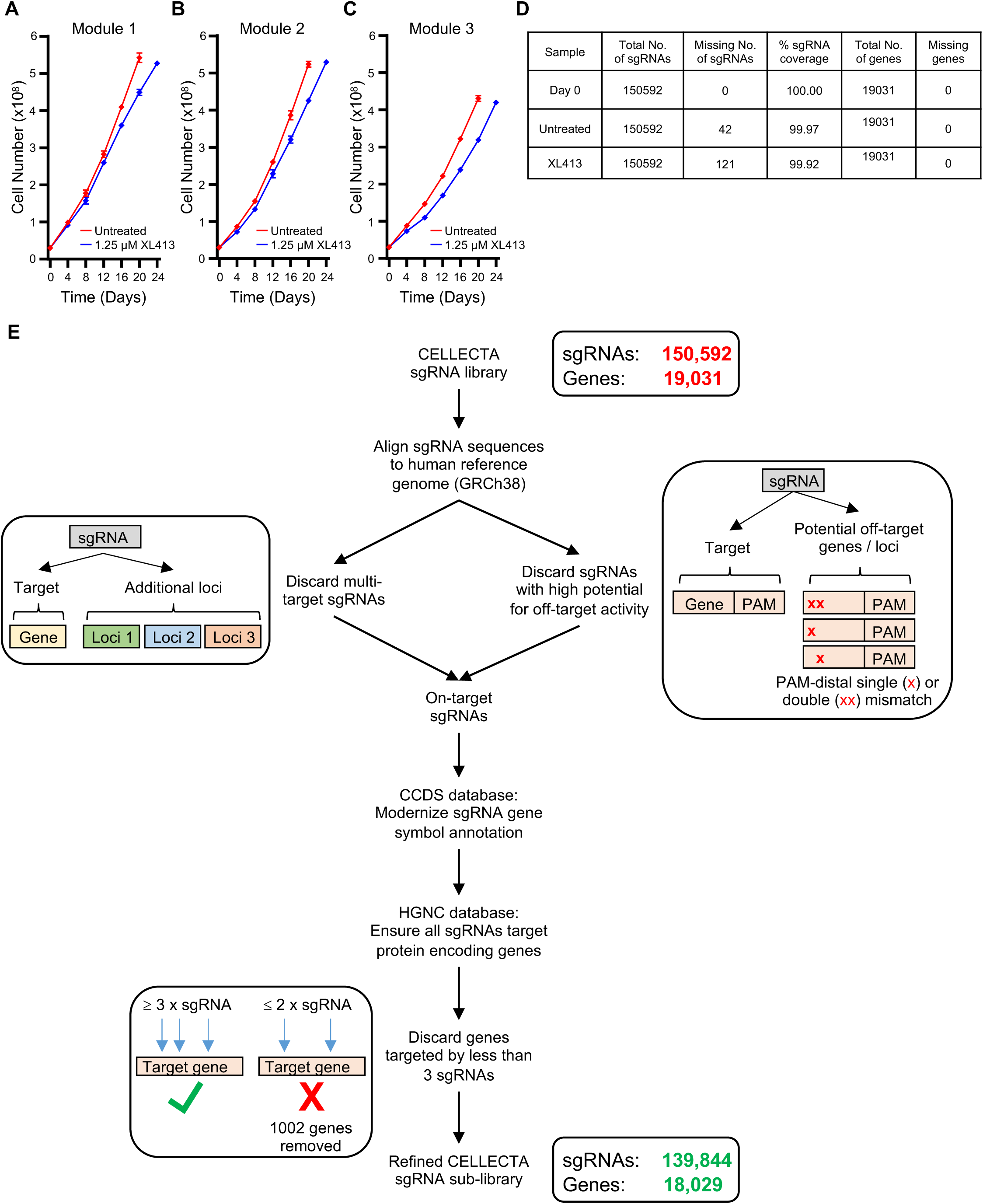
CRISPR/Cas9 screen: Cell proliferation curves, summary of sgRNA coverage, and *in silico* sgRNA library refining scheme. **(A - C)** Proliferation curves of MCF10A EditR cells expressing sgRNAs from CRISPR library modules 1, 2 and 3 in the absence (untreated, red line) or presence of 1.25 µM XL413 (blue line). Data is from three independent counts of each sample and the mean ± SD is plotted. **(D)** DNA extracted from samples of the CRISPR/Cas9 screen, which were harvested on day 0 and after 20 or 24 days of growth in the absence (untreated) or presence of XL413, respectively, was analysed by next generation sequencing. Sequence data was mapped to the Cellecta sgRNA reference library using MAGeCK. The table summarises the number of missing sgRNAs, the calculated % sgRNA coverage and the number of missing genes. **(E)** Schematic of the *in silico* pipeline used to eliminate ‘poor quality’ sgRNAs, update gene symbol nomenclature and, following library refinement, to disregard genes targeted by less than 3 sgRNAs (described in the methods detail). The resulting virtual sgRNA sub-library now consisting of 139,844 sgRNAs targeting 18,029 protein coding gens was subsequently used for high confidence hit calling.

**Supplemental Figure 2:**
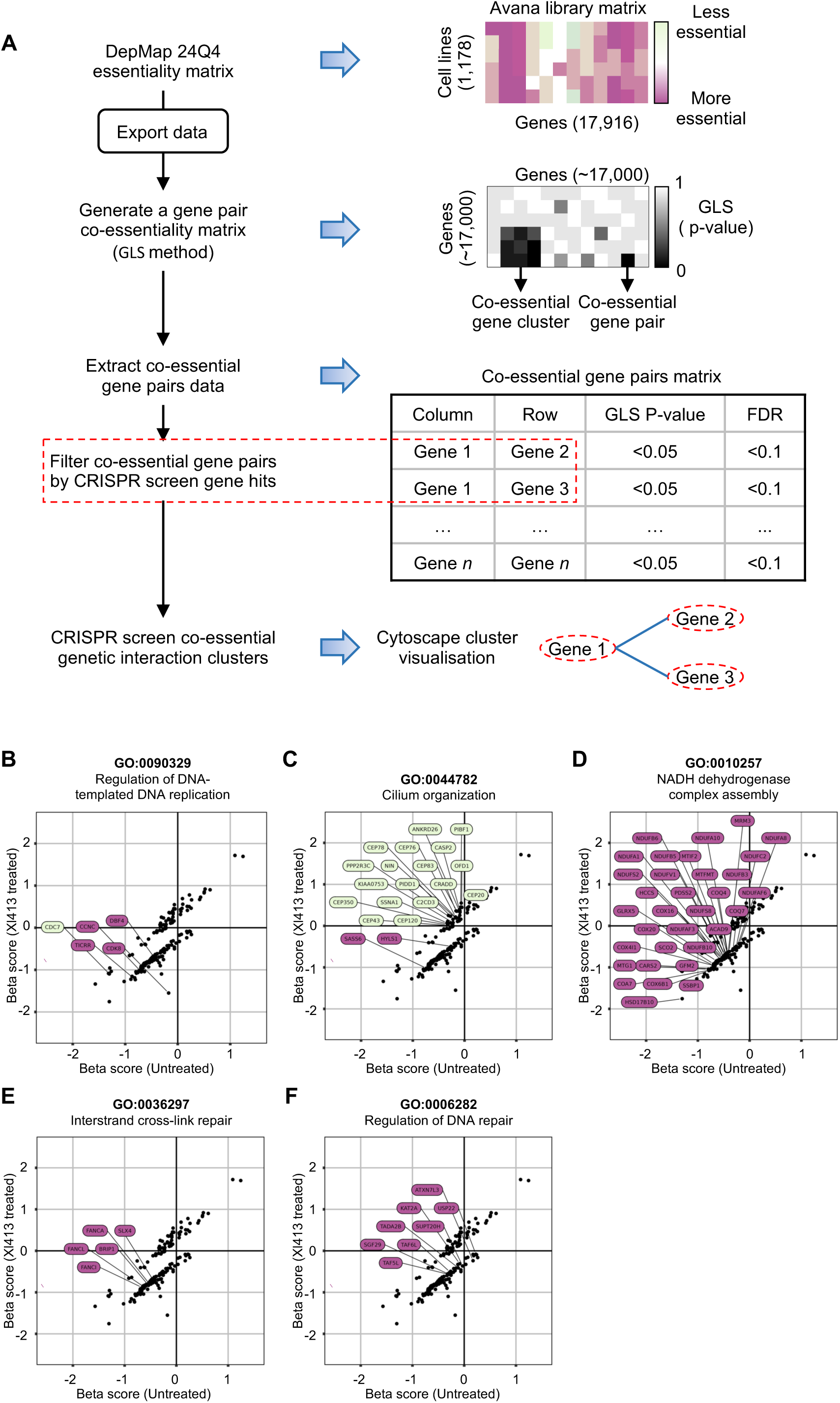
Schematic of *in silico* workflow to visualise co-essential genetic interaction clusters and beta scores of genes in key networks. **(A)** Representation of the *in silico* pipeline used for visualisation of CRISPR screen gene hits as clusters, based on co-essential genetic interactions (described in methods). **(B - F)** MAGeCKFlute Beta scores of all the CRISPR screen hits for the XL413-treated versus the untreated branch are plotted and genes from unique clusters are highlighted in each panel alongside a gene ontology identifier and term. Genes that when knocked-out are associated with negative ΔB score, thus potentially enhancing the anti-proliferative effects of CDC7 are in magenta, while the ones associated with positive ΔB score are in green.

**Supplemental Figure 3 - related to Figure 4:**
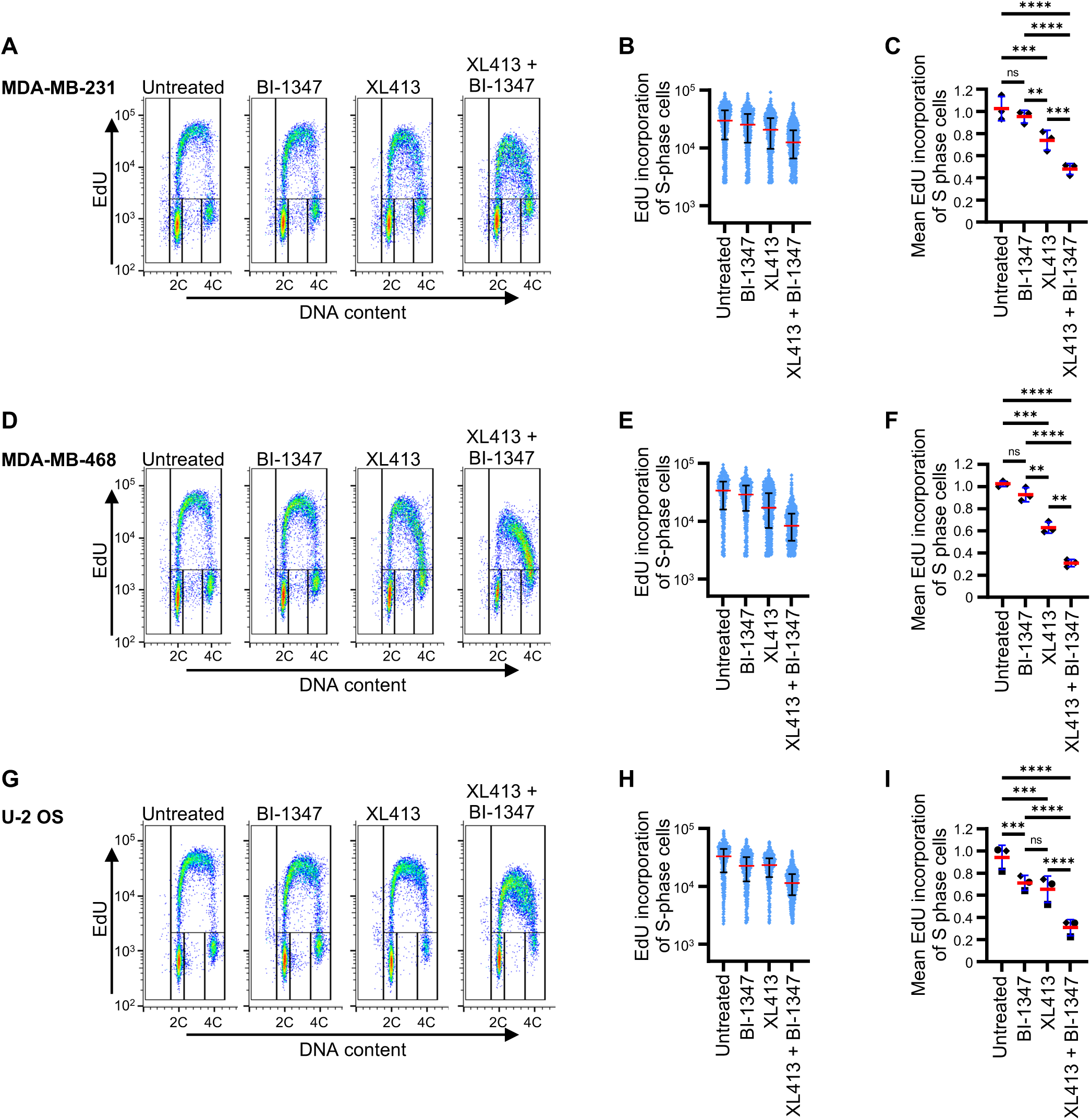
CDK8 and CDC7 inhibitors cooperate to restrict DNA synthesis in multiple cell lines MDA-MB-231. **(A - C)**, MDA-MB-468 **(D - F)**, and U-2 OS **(G - I)** were either mock treated or treated with 31 nM BI-1347, in the presence or absence of 1.25 µM XL413 for 24 hours. **(A, D and G)** Cells were labelled with EdU for last 30 minutes and DNA synthesis, DNA content and cell cycle distribution were assessed by flow cytometry. **(B, E and H)** Fluorescence intensity, proportional to EdU incorporation, in individual S phase cells. Red and blue lines indicate the median and the interquartile range extending from the 25^th^ to the 75^th^ percentile for each condition. **(C, F and I)** Mean fluorescence intensity in all S phase cells from three independent experiments is expressed as a ratio relative to the untreated condition. Red and blue line shows the mean ± SD. Statistical analysis: Two-way ANOVA with Tukey’s multiple comparison post-test; ns, not significant; *P< 0.05; **P< 0.01; ***P< 0.001; ****P< 0.0001.

**Supplemental Figure 4 - related to Figure 7:**
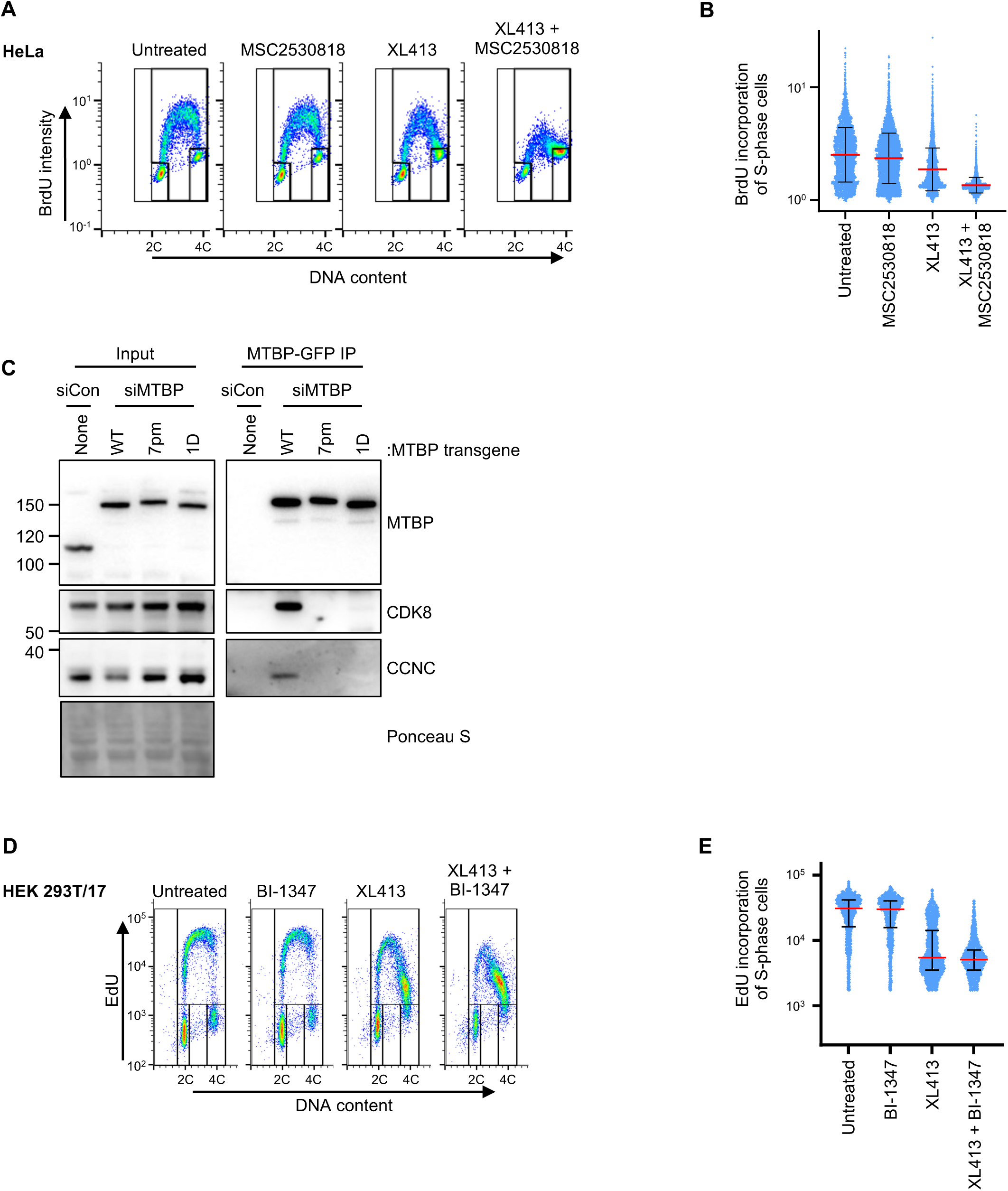
MTBP mutants are defective in binding CDK8-CCNC. **(A)** HeLa Flip-In T-Rex cells were either mock treated or treated with 2 µM MSC2530818 (CDK8 inhibitor), in the presence or absence of 2.5 µM XL413 for 24 hours. Cells were labelled with BrdU for last 1 hour and DNA synthesis, DNA content and cell cycle distribution were assessed by flow cytometry. **(B)** Fluorescence intensity, proportional to BrdU incorporation, in individual S phase cells. Red and blue lines indicate the median and the interquartile range extending from the 25^th^ to the 75^th^ percentile for each condition. **(C)** HEK 293T/17 cells were transfected with C-terminally tagged (3xFlag-TEV2-GFP) MTBP wild type (WT), MTBP-Cdk8-non-binding: seven point mutant (7pm); or single point mutant (1D), or left untransfected. Anti-GFP immunoprecipitations from native cell lysates were performed and analysed by immunoblotting for the indicated proteins. **(D)** HEK 293T/17 cells were either mock treated or treated with 31 nM BI-1347, in the presence or absence of 10 µM XL413 for 24 hours. Cells were labelled with EdU for last 30 minutes and DNA synthesis, DNA content and cell cycle distribution were assessed by flow cytometry. **(E)** Fluorescence intensity, proportional to EdU incorporation, in individual S phase cells. Red and blue lines indicate the median and the interquartile range extending from the 25^th^ to the 75^th^ percentile for each condition.

## SUPPLEMENTAL TABLE LEGEND

Supplementary table 1 related to Figure 1: **Summary of CRISPR/Cas9 screen analysis using MAGeCKFLUTE pipeline.** DNA extracted from samples of modules 1, 2 and 3 (M1, M2 and M3) of the CRISPR/Cas9 screen, which were harvested on day 0 and after growth for 20 days in the absence (untreated) or 24 days in the presence of XL413, was analysed by next generation sequencing. For each module sequence and read count data were mapped to the refined CELLECTA sgRNA sub-library and analysed using MAGeCKFlute pipeline. MAGeCK MLE was used to assign beta scores to genes based on changes of sgRNA abundance from day 0 to day 20 and day 24 in the untreated and XL413 treated samples respectively. Differential beta score (ΔB = XL413-treated Beta score - untreated Beta score) was then used to rank and identify potential gene candidates. The table summarises: the rank order, by differential beta score for genes associated with modules 1, 2 and 3 alongside the number of sgRNAs targeting each gene, the beta score associated with each gene in the XL413-treated and untreated samples, Uniprot identifier (Uniprot_ID) and protein name.

## Notes

### Summary of Updates

Minor changes/edits in the text for improved clarity; ORCID for authors updated.

